# Systematic differences in protein stability underlie species-specific developmental tempo

**DOI:** 10.1101/2024.06.07.597977

**Authors:** Mitsuhiro Matsuda, Henrik M. Hammarén, Jorge Lázaro, Mikhail M. Savitski, Miki Ebisuya

## Abstract

Human embryonic development proceeds more slowly than in mice. The segmentation clock offers a tractable model to study interspecies differences in developmental tempo, as its oscillation period in human induced presomitic mesoderm (iPSM) cells is approximately twice that of mouse. While the core clock gene HES7 exhibits slower protein degradation in human cells, it remains unclear whether such cross-species differences in protein stability reflect a general principle. Here, we perform a dynamic SILAC-based proteomic analysis of ∼5,000 proteins in human and mouse iPSM, and we uncover a pervasive trend of slower protein degradation in human cells, regardless of subcellular localization or degradation pathways. Moreover, inhibition of glycolysis in mouse iPSM phenocopies the human protein stability profile, and modulation of protein stability alters the tempo of both the segmentation clock and cellular differentiation. Together, our findings establish protein stability, with pervasive differences across species, as a key mediator linking metabolism to developmental tempo.

## Introduction

The molecular and cellular mechanisms of embryonic development are largely conserved across mammals, yet the pace of development varies considerably between species^1^. For example, human embryogenesis from fertilization to organogenesis takes ∼60 days, whereas in mice it takes only ∼15 days^2^. Neural differentiation and maturation also take longer in humans than in mice, even though they follow comparable gene expression programs^3–5^. Such interspecies differences in developmental tempo, despite the conserved order and proportions of events, are referred to as developmental “allochrony”^3,6–8^.

One of the most well-studied model systems for allochrony is the segmentation clock, the oscillatory gene expression during embryogenesis that regulates the timing of sequential body segment formation^9^. The oscillation period of the segmentation clock is notably species-specific^6,9–13^: around 5 h in humans, 4 h in rhinoceroses, 2.5 h in mice, and 30 min in zebrafish.

The segmentation clock can be recapitulated in vitro by inducing presomitic mesoderm (PSM) from pluripotent stem cells^14–17^. This iPSM approach enables access to otherwise inaccessible species, such as human and rhinoceros, and allows quantitative cross-species comparisons^6,8,12,18^. Importantly, the ∼5 h and ∼2.5 h periods of the human and mouse segmentation clocks have been reproduced in vitro across different protocols and cell lines, consistent with the pace of in vivo somitogenesis^14–17^. These robust, cell-intrinsic properties make the in vitro segmentation clock a powerful model for studying allochrony.

Using this in vitro platform, several molecular mechanisms underlying species-specific segmentation clock tempo have been investigated. In particular, biochemical reactions of the core clock gene HES7 are slower in human iPSM cells than in mouse iPSM cells, including protein and mRNA degradation as well as gene expression delays^6,12,19^. These differences in HES7 kinetics account for the ∼2-fold longer oscillation period in human cells. In addition, cellular metabolism has emerged as another mechanism^7,20,21^: human iPSM cells display lower metabolic activities compared with mouse iPSM, and their experimental manipulation alters the segmentation clock period^18,22^. Together, these findings suggest that both gene-specific biochemical kinetics and metabolic activities may underlie species-specific segmentation clock tempo.

Despite these advances, several fundamental questions remain unresolved. First, previous measurements of degradation rates and expression delays have focused on a handful of genes such as HES7^6,12,19^, leaving it unclear whether species-specific kinetics is an exceptional feature of a few regulators or a general property of many genes. A systematic comparison of biochemical kinetics between human and mouse iPSM cells has not yet been conducted. Such a comprehensive analysis could provide mechanistic insights and reveal, for example, whether proteins with species-specific degradation rates share common characteristics in their localization or degradation pathways.

Second, while several metabolic activities have been proposed as determinants of developmental tempo^7,18,20–23^, their connection to biochemical kinetics is still poorly understood. Mitochondrial electron transport chain (ETC) activity modulates the segmentation clock period by altering translation and intron processing, but not protein degradation^18,22^. By contrast, glycolysis inhibition decelerates HES7 protein degradation^22^, yet whether this metabolic regulation of protein stability represents a general, proteome-wide principle and through what mechanisms remains unclear.

Third, correlations between protein stability and diverse aspects of biological tempo have been reported, including the period of the segmentation clock, the pace of neural differentiation, and even organismal lifespan^3,6,12,24,25^. However, a causal link has not been established: whether direct perturbation of protein stability is sufficient to alter developmental tempo remains an open question.

In this study, we address these three gaps by systematically quantifying protein degradation rates in human and mouse iPSM, examining the proteome-wide impact of glycolytic activity, and perturbing protein stability to test its causal role in developmental tempo.

## Results

### SILAC-proteomics comparison of human and mouse iPSM cells

To systematically compare protein stability between human and mouse segmentation clocks, we performed dynamic SILAC (stable isotope labeling of amino acids in cell culture) and quantitative mass spectrometry^26–28^. Human and mouse iPSM cells were derived through in vitro differentiation of human induced pluripotent stem cells (iPSCs) and mouse epiblast stem cells (EpiSCs), respectively (Fig. 1a)^6,12,16^. For both species, we employed HES7-knockout stem cell lines to suppress the segmentation clocks and prevent protein levels from oscillating during sampling. As HES7 is not essential for PSM differentiation, HES7-knockout cells undergo normal differentiation and still show increased HES7 promoter activity upon iPSM induction^16^. For the labeling of newly synthesized proteins, iPSM cells were cultured in SILAC medium containing heavy arginine and lysine and collected at multiple time points between 0.5 and 12 h (Fig. 1a). LC-MS/MS (liquid chromatography-coupled tandem mass spectrometry) analysis and protein copy number estimation^29^ suggested that protein copy numbers per cell were comparable between human and mouse iPSM cells (Supplementary Fig. 1). The detected proteins included PSM marker genes such as TBX6 and DLL1, confirming normal iPSM induction from HES7-knockout cells.

**Figure 1.**
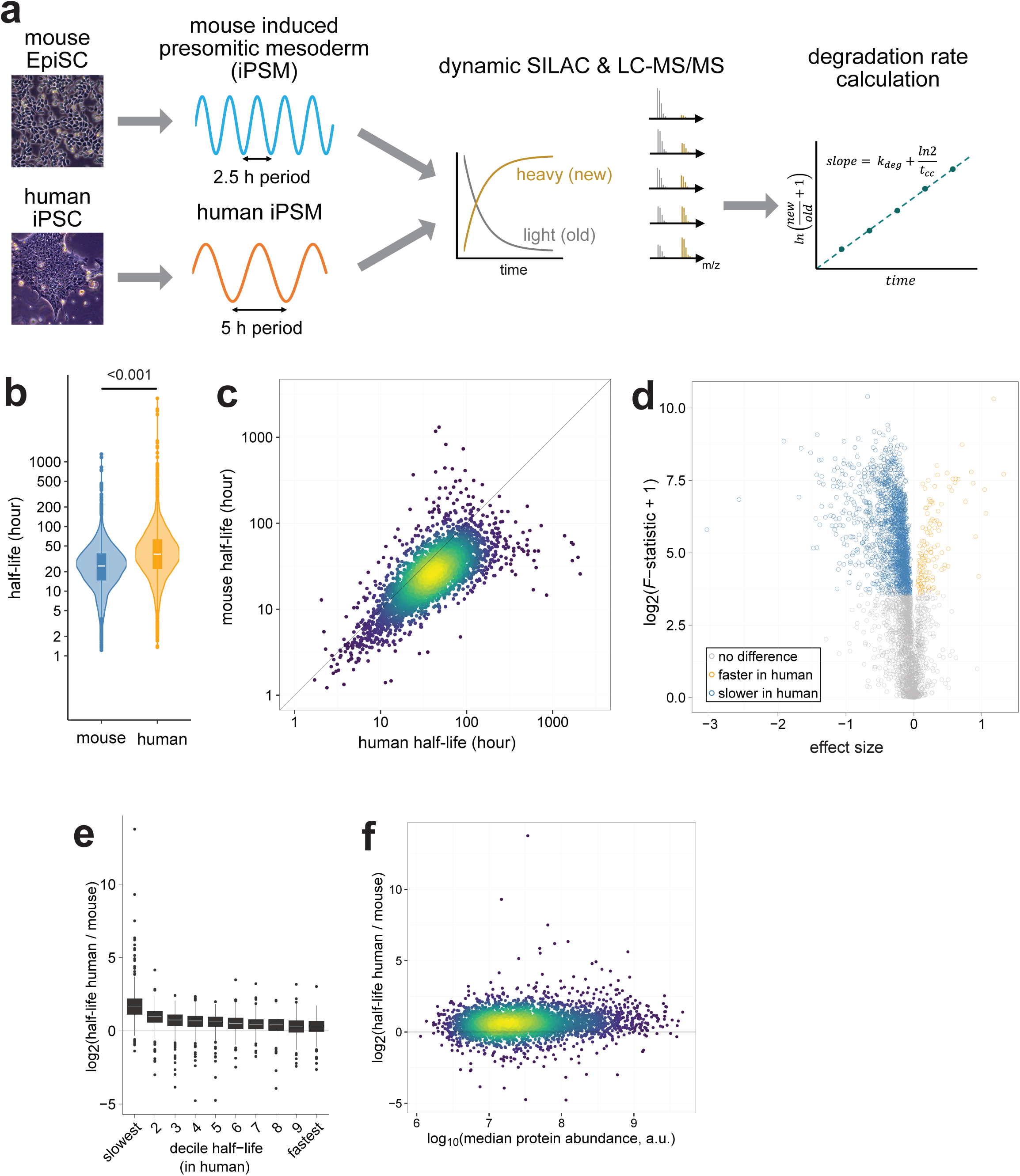
Pervasively slow protein degradation in human iPSM compared with mouse. **a,** Experimental workflow of dynamic SILAC-based proteomics. Human and mouse iPSM cells were derived from human induced pluripotent stem cells (iPSCs) and mouse epiblast stem cells (EpiSCs), respectively, and treated with the heavy SILAC medium from 0.5 h to 12 h for the labeling of newly synthesized proteins. Following LC-MS/MS analysis and peptide quantification, the degradation rate of each protein was calculated based on the ratio of the heavy (new) to light (old) proteins at multiple points. The effect of protein dilution through cell division was compensated. **b,** The distribution and median of protein half-lives in mouse and human iPSM. N = 4835 (mouse), 4612 (human). P-value is from two-sided Mann-Whitney U test. **c,** Scatter plot comparing protein half-lives measured in human and mouse iPSM. N = 3630. **d,** Volcano plot showing proteins with significantly longer (blue) or shorter (orange) half-lives in human cells compared with mouse. The x axis shows the difference in fit between the two species signed by the difference in slope: sign(Δslope)*√(RSS0−RSS1), where RSS0 and RSS1 are the residual sums of squares of the fits for the H0 and H1 model respectively. The y axis shows the F-statistic, a statistical measure of significance, where large values denote greater significance. Proteins with significantly different half-lives (p-value adjusted for multiple testing <= 0.001) are colored. N = 3630. See Methods as well as Supplementary Fig. 2 for more details on the comparative fitting approach. **e,** Fold changes between human and mouse half-lives that were classified according to the human half-lives. **f,** Fold changes between human and mouse half-lives were evenly distributed across the median protein abundance of human and mouse. N = 3630. a.u.: arbitrary units. **b,e,** Box plots consist of median line, box: upper and lower quartiles, whiskers: 1.5 times interquartile range, and points: outliers.

We determined the ratio of newly synthesized (i.e., labeled with heavy amino acids) proteins to preexisting (i.e., light) proteins and calculated protein degradation rates as previously described^27,28^ (Supplementary Fig. 2, see also Methods). Note that the degradation rates were corrected by the protein dilution rate resulting from cell division^27^. The cell cycle time was estimated by assuming that the top 1% of the slowest turning-over proteins decay exclusively through dilution, not degradation. The estimated cell cycle time in human iPSM was longer than that in mouse iPSM (Supplementary Fig. 3a), indicating that protein dilution was also slower in human cells as previously reported^18^. For ease of interpretation, the cell cycle-corrected degradation rates were transformed into half-lives, which are used throughout the rest of the study. To ensure robust estimation of half-lives, human and mouse iPSM samples were measured in three independent replicates. Despite slight variations in PSM differentiation protocols, SILAC pulse timings, and measurement instruments, the proteomic profiles were consistent within samples of the same species (Supplementary Fig. 3b,c), and the estimated half-lives exhibited strong reproducibility (Supplementary Fig. 3d). For the final dataset, we included all gene names with quantified half-lives in at least two out of the three replicates for each species, resulting in half-lives for 4835 mouse and 4612 human gene names, with 3630 shared across species (Fig. 1b,c; Supplementary Table 1).

### Protein degradation in human cells is pervasively slower than in mouse

The distribution of the measured half-lives revealed a consistent pattern of slower degradation in human iPSM compared with mouse iPSM (Fig. 1c). The median half-life was 37 h for human and 24 h for mouse (Fig. 1b). The overall 1.5-fold change between the two species was comparable to the 2-fold change between the human and mouse HES7 degradation rates previously reported^6^. To analyze individual cases of significantly different degradation rates, we used an F-statistic approach^30,31^ (Supplementary Fig. 2e,f, see also Methods). Among the 3630 gene names, 1753 showed significantly slower degradation in human cells while only 126 showed significantly faster degradation in human cells (Fig. 1d). The interspecies difference was more pronounced for slowly turning-over proteins than for rapidly turning-over proteins, even though proteins of all half-life decile ranks showed slower degradation in human cells (Fig. 1e). The interspecies difference in half-lives was independent of protein expression level (Fig. 1f; Supplementary Fig. 3e). These results demonstrated that protein degradation in human iPSM was pervasively slower than in mouse iPSM.

### Human cells show slower protein turnover across compartments and degradation pathways

To characterize the proteins that exhibited significant differences in half-lives between species, we performed gene ontology (GO) analyses. No GO term in molecular function (MF) or biological process (BP) was significantly enriched, likely reflecting pervasive interspecies differences across all functional categories. Protein classification according to the cellular component (CC) GO terms again revealed the pervasive trend of slower degradation in human iPSM than in mouse iPSM, regardless of subcellular localization (Fig. 2a). However, CC GO terms associated with the mitochondrion, endoplasmic reticulum (ER), and membranes were significantly enriched in the proteins that exhibited slower degradation in human cells (Fig. 2b). Consistent with this observation, proteins localized in the mitochondrion and ER showed significantly larger fold changes between human and mouse half-lives compared with the average change (Fig. 2a). By contrast, proteins in the nucleolus showed a smaller fold change, even though their degradation was still slower in human cells (Fig. 2a). These results suggested that proteins, irrespective of localization, turned over more slowly in human cells than in mouse cells and that membrane proteins tended to show larger interspecies differences.

**Figure 2.**
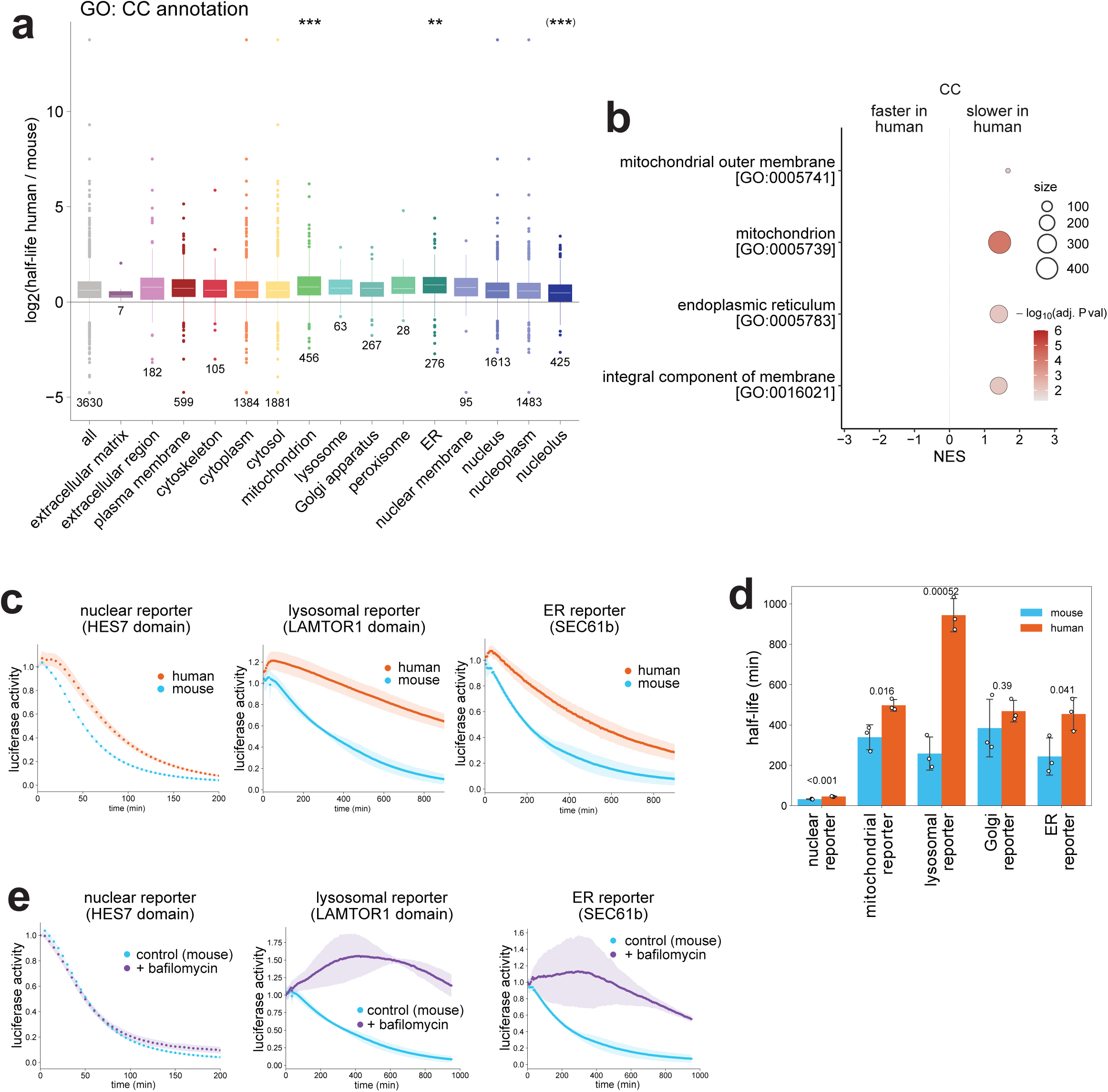
Slower protein degradation in human iPSM across compartments and degradation pathways. **a,** Fold changes between human and mouse half-lives classified according to the Gene Ontology (GO) cellular component (CC) terms. Box plots consist of median line, box: upper and lower quartiles, whiskers: 1.5 times interquartile range, and points: outliers. P-values are from two-sided t test of each annotation group against the reference group of all proteins adjusted for multiple testing. **: p < 0.01, ***: p < 0.001. Cases with p > 0.05 are not shown. Note that proteins localized in the mitochondrion and ER showed significantly larger fold changes in half-lives compared to the average fold change (normal asterisks), while proteins in the nucleolus showed a significantly smaller fold change compared to the average despite still a longer half-life in human than mouse (asterisk in parentheses). **b,** GO CC terms significantly enriched in proteins that showed longer half-lives in human iPSM compared with mouse. P-values are probability estimates that the given enrichment/depletion of the particular gene sets are non-random (two-sided, see fgsea^55^) adjusted for multiple testing using the Benjamini-Hochberg correction. GO terms with adjusted p-value < 0.01 are shown. NES: Normalized Enrichment Score. **c,** Degradation assays of localization reporters in human and mouse iPSM. The transcription of the localization reporters fused with Nano luciferase (Nluc) was halted by doxycycline (Dox) at time 0, and the decay of the luciferase activity was monitored. **d,** Protein half-lives of the localization reporters estimated from c and Supplementary Fig. 4,5. Graph indicates mean ± SD (N = 3). P-values are from two-sided t test. **e,** Degradation assays of localization reporters in the presence of a lysosomal inhibitor bafilomycin A1. Mouse iPSM was treated with 80 nM bafilomycin from 4 h before Dox addition. Control (mouse) data are the same as c. **c,e,** Shading indicates mean ± SD (N = 3).

To verify the results with an independent degradation assay, we created a series of luciferase-fusion reporters representing different subcellular localizations (Supplementary Fig. 4). The nuclear reporter comprised the DNA-binding domain of HES7 and luciferase. The mitochondrial, lysosomal, Golgi, and ER reporters employed the localization domains or full-length sequences of ATP5MC1, LAMTOR1, NOS3, and SEC61B, respectively (Supplementary Fig. 4a). We measured the degradation rates of the localization reporters by halting their transcription and monitoring the decrease in the luciferase signal. All the localization reporters except for the Golgi reporter displayed significantly slower protein degradation in human iPSM compared with mouse iPSM (Fig. 2c,d; Supplementary Fig. 4b,5), consistent with the SILAC-proteomics results.

A potential commonality among the CC GO terms that showed the most significant interspecies differences (i.e., membranes, ER, and mitochondrion) is that the proteins in those localizations are mostly degraded through lysosomes^32,33^. Intracellular proteins are degraded through two main pathways: the ubiquitin-proteasome and autophagy-lysosome systems. Degradation through the ubiquitin-proteasome system has been shown to exhibit interspecies differences: the bulk proteasome activity is lower in human iPSM than in mouse iPSM, and HES7, a control protein that displays the interspecies difference, is ubiquitinated and degraded through proteasomes^12,18,34^. By contrast, it has been unclear whether protein degradation through the autophagy-lysosome system has interspecies differences. To address this question, we inhibited autophagy and lysosomal activities with bafilomycin A1, a vacuolar-type H+-ATPase blocker^35^. Treatment of mouse iPSM with bafilomycin inhibited the degradation of the mitochondrial, lysosomal, Golgi, and ER reporters, indicating that these degradations were indeed mediated through the autophagy-lysosome system (Fig. 2e; Supplementary Fig. 4c). By contrast, bafilomycin treatment did not affect the degradation of the nuclear reporter, as HES7 degradation is mediated through the ubiquitin-proteasome system^34^ (Fig. 2e). Consistent with this, the degradation of the nuclear reporter was inhibited by treatment with a proteasome inhibitor MG-132 (Supplementary Fig. 4d). These results collectively demonstrated that protein degradation was pervasively slower in human cells than in mouse cells, not only through the ubiquitin-proteasome system but also the autophagy-lysosome system. The greater interspecies differences observed in lysosomal substrate stability may reflect the higher variability of the autophagy-lysosome system across mammals compared with the more conserved ubiquitin-proteasome system^36^.

### Glycolysis inhibition decelerates the segmentation clock and protein degradation

We next investigated the effect of metabolism on the protein stability profile. While both glycolytic and ETC activities contribute to the segmentation clock and somitogenesis^12,18,37–41^, we recently found that glycolysis inhibition decelerates HES7 protein degradation whereas ETC inhibition extends HES7 intron processing delay^22^. Given the emphasis of this study on protein stability, we focused on the glycolysis inhibitor 2-deoxy-D-glucose (2DG). As reported previously^22^, treatment of mouse iPSM with 2DG decelerated the segmentation clock in a dose-dependent manner (Supplementary Fig. 6a,b): the oscillation period of mouse iPSM treated with 10 mM 2DG was 191 ± 1 min (mean ± SD), which was between the untreated mouse iPSM period (147 ± 4 min) and the reported human iPSM period (322 min)^6^. The 2DG treatment also decelerated HES7 protein degradation in a dose-dependent manner^22^ (Supplementary Fig. 6c,d): the HES7 half-life in mouse iPSM treated with 10 mM 2DG was 34 ± 3 min (mean ± SD), which was closer to the reported half-life in human iPSM (40 min)^6^ than to untreated mouse iPSM (22 ± 0.8 min). The HES7 period extension by 2DG was limited compared with the half-life extension (compare Supplementary Fig. 6b with 6d), likely because 2DG does not effectively influence other important parameters contributing to the period, such as HES7 intron processing delay^22^. We further confirmed that the 2DG treatment decreased glycolytic activity^22^ (Supplementary Fig. 7).

To examine the impact of glycolytic activity on the global protein stability profile, we performed the dynamic SILAC-proteomics with mouse iPSM treated with 10 mM 2DG (hereafter referred to as mouse_2DG). The cell cycle time of the mouse_2DG sample was closer to that of the human sample than the mouse sample (Supplementary Fig. 3a). The distribution of the measured half-lives revealed a consistent pattern of slower degradation in mouse_2DG compared with untreated mouse (Fig. 3a; Supplementary Table 1). Among 3945 gene names, 1359 showed significantly slower degradation in mouse_2DG while only 24 showed significantly faster degradation in untreated mouse (Fig. 3b). The half-life extension by 2DG showed a very slight tendency to be greater for highly abundant proteins (Supplementary Fig. 8). Unlike the human versus mouse sample comparison (Fig. 1e), the mouse_2DG versus mouse comparison did not exhibit larger differences for slowly turning-over proteins (Fig. 3c). Broad CC GO terms, including those associated with the proteasome, nucleolus, and ribosome, were enriched in the proteins that exhibited slower degradation in mouse_2DG (Supplementary Fig. 9a). These results indicated a broad impact of glycolytic activity on protein degradation.

**Figure 3.**
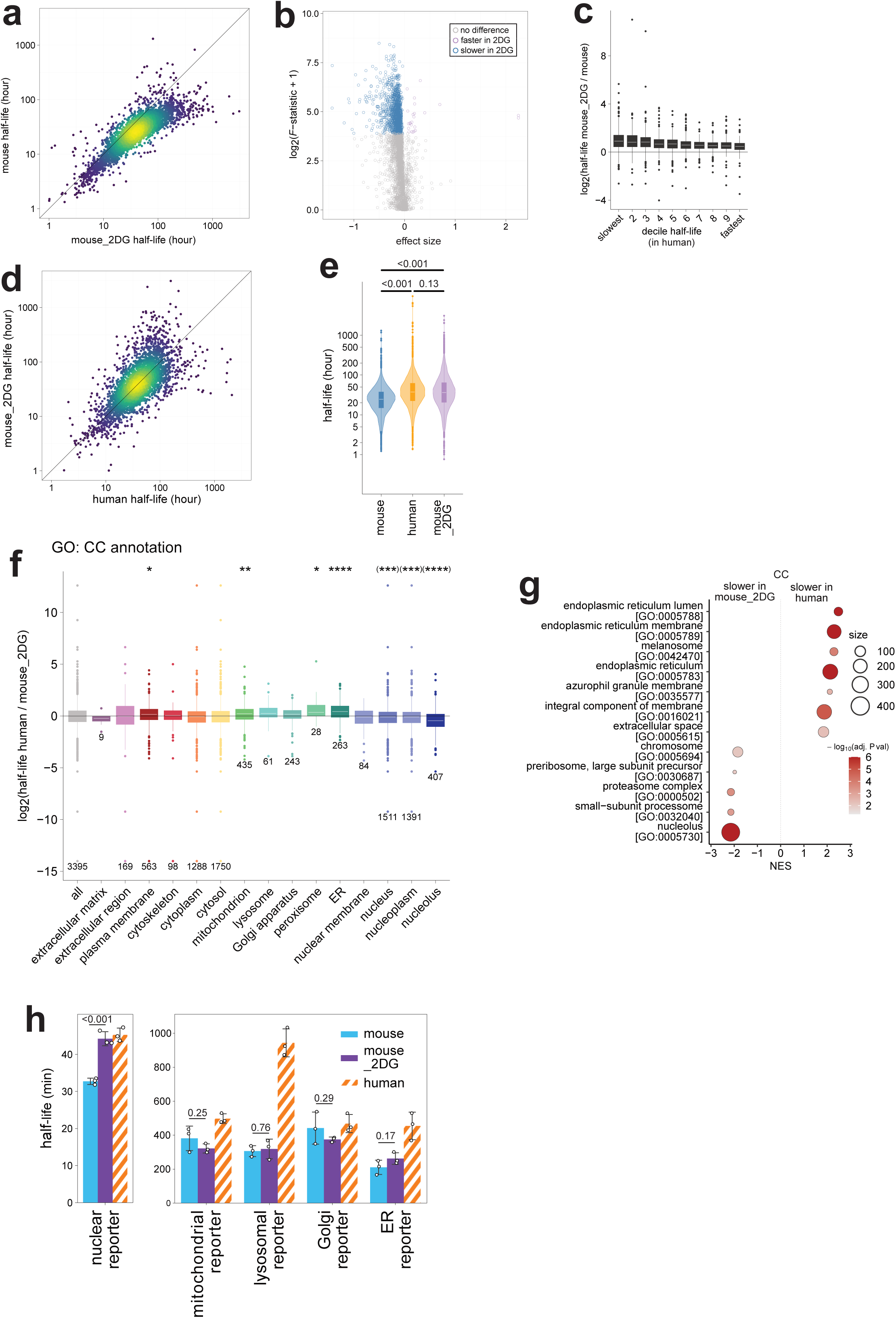
Similarities of protein stability profiles between glycolysis-inhibited mouse iPSM and human. **a,** Scatter plot comparing protein half-lives measured in mouse iPSM treated with 10 mM 2DG (mouse_2DG) and untreated mouse iPSM (mouse). N = 4083. **b,** Volcano plot showing proteins with significantly longer (blue) or shorter (magenta) half-lives in mouse_2DG compared with mouse. The x axis shows the difference in fit between the two samples signed by the difference in slope: sign(Δslope)*√(RSS0−RSS1), where RSS0 and RSS1 are the residual sums of squares of the fits for the H0 and H1 model respectively. The y axis shows the F-statistic. Proteins with significantly different half-lives (p-value adjusted for multiple testing <= 0.001) are colored. N = 3945. See Methods as well as Supplementary Fig. 2 for more details. **c,** Fold changes between mouse_2DG and mouse half-lives that were classified according to the human half-lives, similarly to Fig. 1e. **d,** Scatter plot comparing protein half-lives measured in human iPSM (human) and mouse iPSM treated with 10 mM 2DG (mouse_2DG). N = 3395. **e,** The distribution and median of half-lives in mouse, human, and mouse_2DG. N = 4835 (mouse), 4612 (human), 4517 (mouse_2DG). P-values are from Kruskal-Wallis test followed by pairwise Mann-Whitney U test with Bonferroni correction. **f,** Fold changes between human and mouse_2DG half-lives classified according to the GO CC terms. P-values are from two-sided t test of each annotation group against the reference group of all proteins adjusted for multiple testing. *: p < 0.05, **: p < 0.01, ***: p < 0.001, ****: p < 0.0001. Cases with p > 0.05 are not shown. Note that proteins localized in the plasma membrane, mitochondrion, peroxisome, and ER showed significantly larger fold changes in half-lives compared to the average fold change (normal asterisks), while proteins in the nucleus, nucleoplasm, and nucleolus showed significantly smaller fold changes compared to the average (asterisks in parentheses). **g,** GO CC terms significantly enriched in genes that showed faster or slower degradation in human compared with mouse_2DG. P-values are probability estimates that the given enrichment/depletion of the particular gene sets are non-random (two-sided, see fgsea^55^) adjusted for multiple testing using the Benjamini-Hochberg correction. GO terms with adjusted p-value < 0.01 are shown. **h,** Protein half-lives of the localization reporters in the presence of 2DG estimated from supplementary Fig. 10,11. Human data are from Fig. 2d. Graphs indicate mean ± SD (N = 3). P-values are from two-sided t test. **c,e,f,** Box plots consist of median line, box: upper and lower quartiles, whiskers: 1.5 times interquartile range, and points: outliers.

### Glycolysis inhibition in mouse cells mimics the human protein stability profile

The protein stability profile of the mouse_2DG sample was much more similar to that of the human sample than the mouse sample (Fig. 3d). The median half-life of mouse_2DG was 36 h, closer to the 37 h in human than the 24 h in mouse (Fig. 3e).

However, the impact of 2DG was somewhat localization-dependent (Fig. 3f,g; Supplementary Fig. 9b). Proteins localized in the nucleus, nucleolus, and proteasome complex showed slower degradation in mouse_2DG compared with human. By contrast, proteins in the membranes, mitochondrion, peroxisome, and ER still showed slower degradation in human compared with mouse_2DG. These results suggested that the inhibition of glycolytic activity impacted proteasome-mediated degradation more effectively than lysosome-mediated degradation. Consistent with this hypothesis, 2DG treatment decelerated the proteasome-mediated degradation of the nuclear reporter to the level observed in human, while it did not significantly influence the lysosome-mediated degradation of the mitochondrial, lysosomal, Golgi, and ER reporters (Fig. 3h; Supplementary Fig. 10,11). This difference in degradation pathways may reflect either the protective upregulation of autophagic activity by 2DG^42^ or the stronger evolutionary conservation of the ubiquitin-proteasome system^36^. Nevertheless, the inhibition of glycolytic activity in mouse cells mostly phenocopied the human protein stability profile.

### Protein stability modulates developmental tempo

Having established the relationship between metabolic activity and protein stability, we next investigated the causal link between protein stability and developmental tempo. Although mathematical models have demonstrated that HES7 protein stability contributes to the segmentation clock period^6,43,44^, this prediction has not been experimentally verified, because decelerating HES7 degradation often disrupts clock oscillations^34^. We analyzed mouse Hes7 mutants in which lysine residues were substituted with arginine to prevent ubiquitination^34^. The Hes7 K14R mutant displayed a degradation rate comparable to that of the wild type (WT), whereas the K211R mutant degraded more slowly (Fig. 4a,b). To evaluate the impact of slower Hes7 protein degradation on the segmentation clock, we introduced Hes7 constructs driven by the Hes7 promoter into Hes7-knockout (KO) iPSM cells. While the exogenous Hes7 constructs rescued oscillations in KO cells, the oscillation period restored by the K211R mutant was longer than that of WT (Fig. 4c,d), demonstrating a connection between the Hes7 degradation rate and clock period. As an alternative approach to modulate protein degradation, we treated iPSM cells with proteasome inhibitors. Because mouse cells were highly sensitive to prolonged pharmacological treatment, we used human iPSM cells and applied bortezomib, a proteasome inhibitor with lower toxicity than MG-132. Broad suppression of protein degradation by bortezomib decelerated the segmentation clock in a dose-dependent manner (Fig. 4e,f). These results provided experimental evidence establishing a causal link between protein stability and the segmentation clock period.

**Figure 4.**
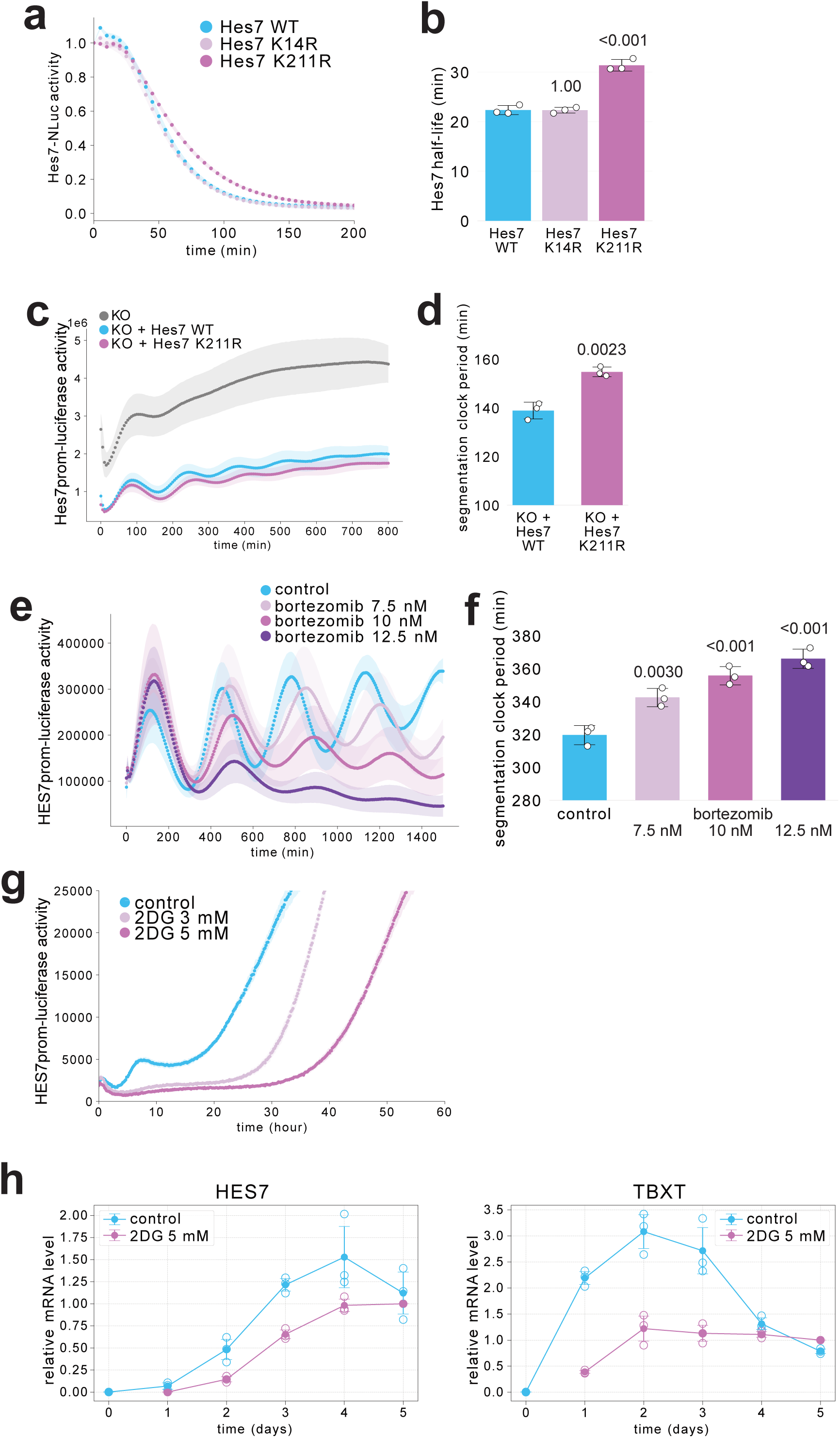
Protein stability-dependent modulation of the segmentation clock and differentiation timing. **a,** Degradation assays of Hes7 variants. Hes7 wild type (WT), K14R, and K211R were fused with Nluc and overexpressed under the rTetOne promoter in mouse iPSM. The transcription of Hes7-Nluc was halted by Dox at time 0, and the decay of the luciferase activity was monitored. **b,** Protein half-lives of Hes7 variants calculated from a. **c,** Oscillation rescue in Hes7-knockout (KO) cells with Hes7 variants. Hes7 WT and K211R were overexpressed under the Hes7 promoter in Hes7-KO mouse iPSM. The oscillatory activity of the Hes7 promoter-luciferase reporter was monitored. **d,** Oscillation periods of Hes7 variants estimated from c. **e,** Dose-dependent effect of a proteasome inhibitor bortezomib on the segmentation clock period. Human iPSM was treated with bortezomib at time 0, and the oscillatory activity of the HES7 promoter-luciferase reporter was monitored. **f,** HES7 oscillation periods estimated from e. **g,** Real-time monitoring of PSM differentiation from human iPSCs in the presence of 2DG. The signal of the HES7 promoter-luciferase reporter was monitored during cellular differentiation, and 2DG treatment delayed the onset of reporter activity. **h**, qRT-PCR analysis of HES7 and TBXT mRNA expression during PSM differentiation from human iPSCs in the presence of 2DG. **a,c,e,g,** Shading indicates mean ± SD (N = 3). **b,d,f,h,** Graphs indicate mean ± SD (N = 3). P-values are from two-sided Dunnett’s test (b,f) or two-sided t test (d) against the indicated controls.

We further explored the impact of protein stability on developmental tempo beyond the segmentation clock, specifically cellular differentiation timing. The onset of HES7 expression was used as a readout for PSM differentiation. Because prolonged bortezomib treatment was too harsh for cells, we instead used 2DG to modulate protein stability. 2DG treatment delayed the initiation of HES7 expression (Fig. 4g), suggesting a slower tempo of PSM differentiation. RT-PCR analysis of PSM differentiation markers (HES7, MSGN1, and TBX6) further revealed delayed expression of all three genes (Fig. 4h; Supplementary Fig. 12), and the transient expression of the NMP marker TBXT was likewise postponed (Fig. 4h). It should be noted, however, that overall expression levels in 2DG-treated cells were lower than in controls, suggesting potential side effects of glycolysis inhibition. Together, these results identified protein stability as a key modulator of developmental tempo, including both the segmentation clock period and cellular differentiation timing.

## Discussion

In this study, we demonstrated that protein degradation is pervasively ∼1.5-fold slower in human iPSM than in mouse iPSM. We further established a mechanistic link between glycolytic activity, global protein stability, and developmental tempo.

Previous studies have measured biochemical kinetics of a handful of marker genes and reporters associated with species-specific developmental tempo^3,6,12,18,19^. While these targeted measurements are precise and valuable, they carry the risk of misinterpretation due to exceptional genes that deviate from the general trend. In this study, by systematically profiling protein stability in the same cell type across two species, we obtained a comprehensive view that enabled the classification of genes with interspecies differences. Degradation via both the ubiquitin-proteasome and autophagy-lysosome systems, as well as dilution by cell division, were all slower in human iPSM than in mouse iPSM. Membrane proteins degraded through the autophagy-lysosome system exhibited particularly large interspecies differences. Consistent with our findings, a recent proteomics study reported ∼1.5-fold longer protein half-lives in human neural progenitors compared with mouse cells^24^, suggesting that the pervasively slower protein turnover in human cells may be a general principle across cell types. Even outside embryonic development, a comparative proteomics study of fibroblasts from twelve mammalian species revealed a correlation between protein stability and organismal lifespan^25^. Thus, systematic comparisons of biochemical kinetics across diverse cell types and species are a promising approach to understanding interspecies differences in developmental and other biological tempos^8^.

Our study also identified glycolytic activity as a potential driver of pervasive differences in protein stability. The extent to which the slower protein degradation in human cells can be attributed to differential glycolytic activity, however, remains unresolved. Human iPSM cells exhibit lower mass-specific glycolysis rates than mouse iPSM^12,18^, yet glycolysis rates do not consistently correlate with segmentation clock periods or HES7 protein degradation rates across or even within species^12,41^. Moreover, how glycolytic activity modulates protein degradation is still unclear. Because both the ubiquitin-proteasome and autophagy-lysosome systems are multi-step, ATP-dependent processes, reduced ATP production may decelerate rate-limiting degradation steps. However, glycolysis and its inhibitor 2DG also affect other signaling and glycosylation pathways^39,41,45,46^. Notably, glycolysis inhibition more strongly impaired proteasome-mediated than lysosome-mediated degradation, possibly because the ubiquitin-proteasome system is evolutionarily more conserved^36^. Defining the exact contribution of glycolysis to the species-specific protein stability profile therefore remains an important direction for future study.

Beyond establishing a correlation, we demonstrated a causal link between protein stability and developmental tempo, including both the segmentation clock and cellular differentiation. Nonetheless, protein stability is unlikely to act as the sole determinant of developmental tempo. For example, the segmentation clock period is influenced not only by protein degradation rates but also by gene expression delays^6,22,43^, whereas epigenetic modifications modulate various aspects of developmental timing^5,47^. Another challenge is to perturb protein stability without off-target effects, underscoring the need for more targeted and less toxic approaches. Looking ahead, long-term modulation of protein stability combined with systematic cross-species comparisons should provide a powerful framework to uncover how protein stability shapes developmental allochrony and biological tempos throughout the life cycle.

## Methods

### Cell cultures and PSM induction

Human iPSCs (201B7 line, RIKEN BRC #HPS0063)^48^ were maintained on Matrigel-coated dishes in StemFit medium (Ajinomoto) under feeder-free conditions. Cells were passaged in the presence of ROCK inhibitor Y-27632 (10 μM). The use of human iPSCs was approved by the Department de Salut de la Generalitat de Catalunya (Carlos III Program) and by the ethics committee at TU Dresden. For human PSM induction, two protocols were applied: 1-step and 2-step. In the 1-step protocol, 2 × 10□ iPSCs were seeded on Matrigel-coated 35 mm dishes and cultured for three days, followed by replacement of the medium with CDMi^49^ containing SB431542 (10 μM), CHIR99021 (10 μM), DMH1 (2 μM), and bFGF (20 ng/ml). This medium is referred to as SCDF medium. Cells were cultured in SCDF medium for an additional three days. In the 2-step protocol, 2 × 10□ iPSCs were plated on Matrigel-coated 35 mm dishes and cultured for 3-4 days. The medium was then replaced with CDMi supplemented with bFGF (20 ng/ml), CHIR99021 (10 μM), and Activin A (20 ng/ml) for one day, followed by culture in SCDF medium for one more day.

Mouse EpiSCs (RIKEN BRC #AES0204)^50^ were maintained on fibronectin-coated dishes in DMEM/F12-based medium containing 15% Knockout Serum Replacement, Glutamax (2 mM), non-essential amino acids (0.1 mM), β-mercaptoethanol (0.1 mM), Activin A (20 ng/ml), bFGF (10 ng/ml), and IWR-1-endo (2.5 μM). Cells were passaged with Y-27632 (10 μM), and medium was refreshed daily. For mouse PSM induction, 5 × 10□ EpiSCs were seeded on Matrigel-coated 35 mm dishes and cultured in the maintenance medium without IWR-1 for one day, followed by SCDF medium for 30 h.

### DNA constructs and reporter cell lines

The localization reporter constructs for degradation assays are illustrated in Supplementary Fig. 4a. To analyze HES7 protein degradation rates, wild type (WT), K14R, and K211R variants^34^ of mHes7 were generated by PCR-based mutagenesis. For the HES7 oscillation assay, the mHes7 promoter-Firefly luciferase (Fluc)-NLS-PEST-stop-mHes7 (w/o intron) construct^6^ was used. For the HES7 protein degradation assay, the reverse TetOne (rTetOne) promoter-mHes7 (w/o intron)-Nano luciferase (Nluc) construct□ was used.

All promoters and genes were subcloned into the pDONR vector to generate entry clones. These entry clones were recombined with the piggyBac vector^51^ (a gift from K. Woltjen) using Multisite Gateway technology (Invitrogen). The constructs were stably introduced into human iPSCs and mouse EpiSCs by electroporation with a 4D Nucleofector (Lonza) or using Lipofectamine (Invitrogen).

The HES7-knockout human iPSC line was previously described□. The Hes7-knockout mouse EpiSC line was generated using the CRISPR guide constructs previously reported for mouse embryonic stem (ES) cells^6^. Multiple CRISPR guide constructs were transiently introduced into EpiSCs, and the deletion of the Hes7 locus was verified by PCR. The disruption of segmentation clock oscillation was further confirmed in mouse iPSM cells derived from the Hes7-knockout EpiSCs.

Oscillation rescue assays were performed exclusively in Hes7-knockout mouse EpiSCs. Specifically, Hes7 WT and K211R variants were expressed from constructs containing the Hes7 promoter, intron, Nluc, and UTR, introduced via piggyBac vectors. For these rescue experiments, cells were analyzed as bulk populations without clonal selection.

### SILAC samples

HES7-knockout human iPSCs and Hes7-knockout mouse EpiSCs were used to suppress segmentation clock oscillations and thereby avoid periodic fluctuations in protein abundance during SILAC sampling. After induction into iPSM, the culture medium was replaced with heavy SILAC medium consisting of SCDF supplemented with heavy lysine (^13^C_6_^15^N_2_, Silantes) and heavy arginine (^13^C_6_^15^N_4_, Silantes). For SILAC culture, IMDM and Ham’s F-12 for SILAC (Gibco #88367 and #88424), which contain minimal levels of light lysine and arginine, were used to prepare the SCDF base medium. SILAC experiments were performed in three independent series with human iPSCs and three series with mouse EpiSCs. For human iPSM induction, the first and second experiments were carried out using the one-step protocol, whereas the third experiment employed the two-step protocol. All three mouse experiments used the same induction protocol with SCDF medium. In addition, one SILAC experiment was performed with 2DG treatment in mouse iPSM cells: following iPSM induction, cells were treated with 2DG for 24 h to allow steady-state adaptation, and subsequently switched to heavy SILAC medium.

Sampling time points differed across experiments: the first experiments (human and mouse) were sampled at 2, 6, and 12 h; the second at 0.5, 1, 1.5, 2, 3, 6, and 12 h; and the third at 1.5, 3, 6, and 12 h. For the 2DG-treated experiment, cells were sampled at 1.5, 3, 6, and 12 h following steady-state adaptation. In 12-h sampling experiments, the heavy SILAC medium was replaced with fresh medium at 6 h to maintain nutrient availability, pH balance, and overall metabolic stability during prolonged culture. Cells were harvested at each time point and lysed in 50 mM HEPES-NaOH (pH 8.5) containing 1% SDS, protease inhibitor cocktail, and benzonase. Lysates were incubated at 95 °C for 5 min prior to downstream processing.

### Sample preparation for mass spectrometry

Protein concentration was measured using PierceTM BCA protein assay kit (Thermo Fisher Scientific #23225), and 10 µg protein was subjected to sample preparation for MS using a modified SP3 protocol^52^. Briefly, protein samples were precipitated onto Sera-Mag SpeedBeads (GE Healthcare #45152105050250 and #65152105050250) in the presence of 50% EtOH and 2.5% formic acid (FA) for 15 min at room temperature, followed by four washes with 70% ethanol on 0.22 µm filter plates (Millipore #MSGVN22). Proteins were digested on beads with trypsin and Lys-C (5 ng/µl final concentration each) in 90 mM HEPES (pH 8.5), 5 mM chloroacetic acid, and 1.25 mM TCEP overnight at room temperature. Peptides were eluted by centrifugation using 2% DMSO and vacuum-dried. Dry peptides were desalted by loading them on an OASIS HLB µElution plate (Waters #186001828BA), washing twice with 0.05% FA, eluting with 80% acetonitrile (ACN), 0.05% FA, and vacuum-drying. Dried peptides were taken up in 20 mM ammonium formate (pH 10) and prefractionated into 12 fractions on an Ultimate 3000 (Dionex) HPLC using high-pH reversed-phase chromatography (running buffer A: 20 mM ammonium formate pH 10; elution buffer B: ACN) on an X-bridge column (2.1 × 10 mm, C18, 3.5 µm, Waters). Prefractionated peptides were vacuum-dried.

### LC-MS/MS analysis

For LC-MS/MS analysis, peptides were reconstituted in 0.1% FA and 4% ACN and analyzed by nanoLC-MS/MS. As different MS methods were used depending on the instrument availability, sample consistency and reproducibility were checked (Supplementary Fig. 3b-d).

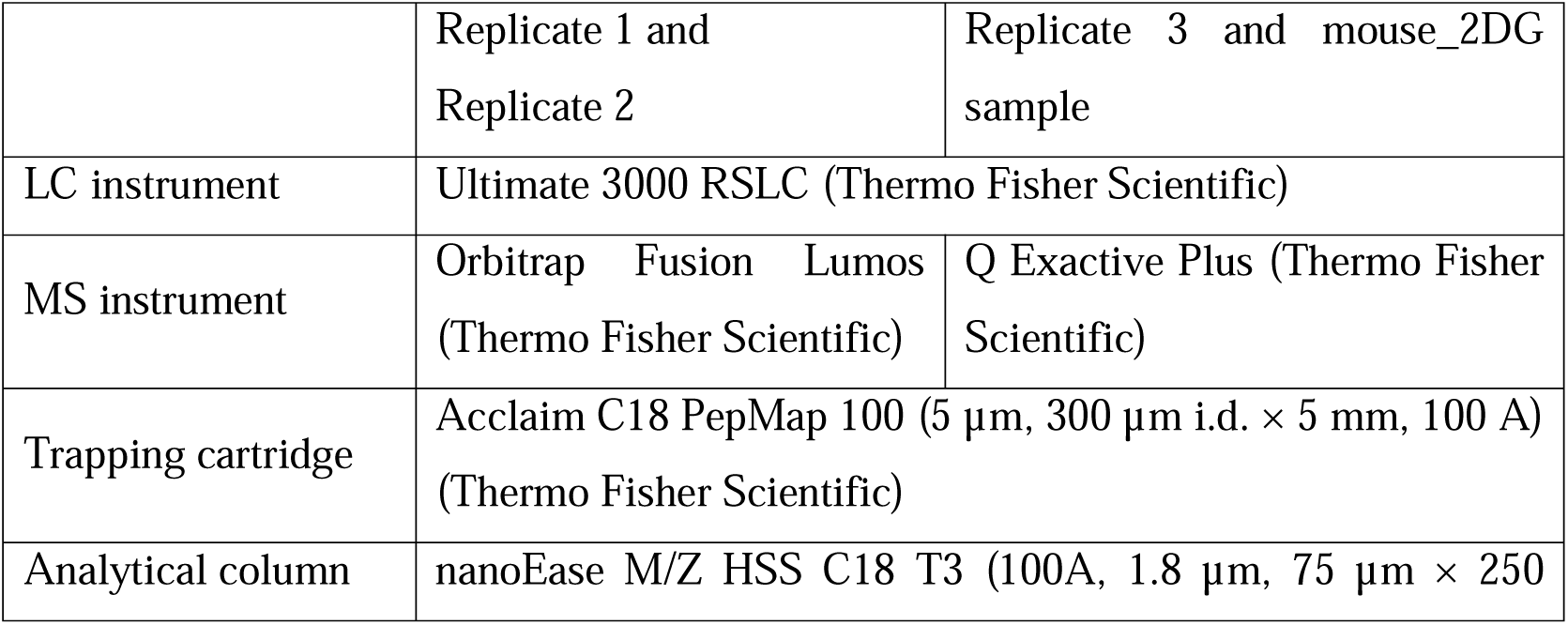
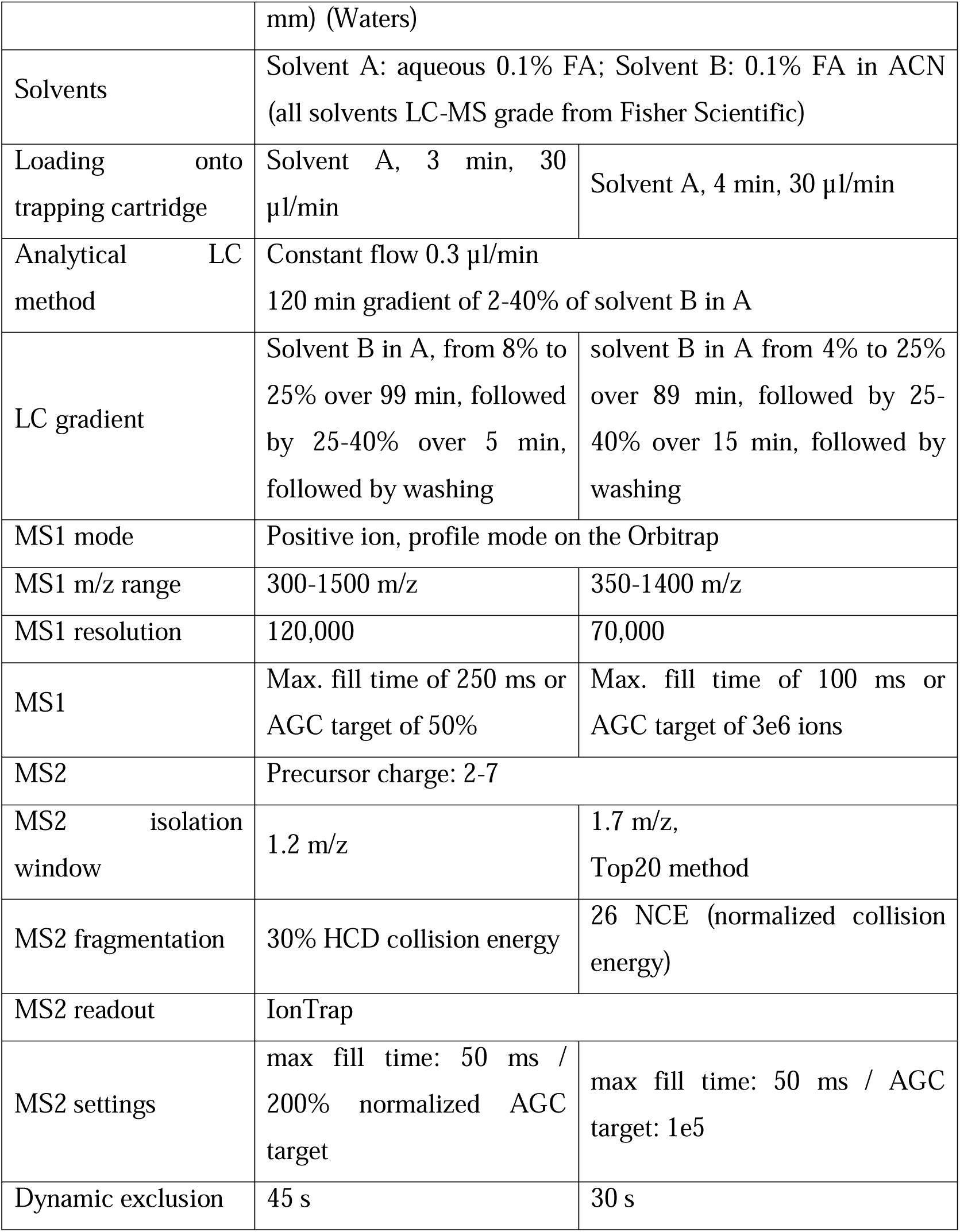

|  |  |  |
| --- | --- | --- |
|  | Replicate 1 and<br>Replicate 2 | Replicate 3 and mouse_2DG<br>sample |
| LC instrument | Ultimate 3000 RSLC (Thermo Fisher Scientific) |  |
| MS instrument | Orbitrap Fusion Lumos<br>(Thermo Fisher Scientific) | Q Exactive Plus (Thermo Fisher<br>Scientific) |
| Trapping cartridge | Acclaim C18 PepMap 100 (5 µm, 300 µm i.d. × 5 mm, 100 Å)<br>(Thermo Fisher Scientific) |  |
| Analytical column | nanoEase M/Z HSS C18 T3 (100Å, 1.8 µm, 75 µm × 250 |  |
|  | mm) (Waters) |  |
| Solvents | Solvent A: aqueous 0.1% FA; Solvent B: 0.1% FA in ACN<br>(all solvents LC-MS grade from Fisher Scientific) |  |
| Loading onto trapping cartridge | Solvent A, 3 min, 30 μl/min | Solvent A, 4 min, 30 μl/min |
| Analytical LC method | Constant flow 0.3 μl/min<br>120 min gradient of 2-40% of solvent B in A | |
| LC gradient | Solvent B in A, from 8% to 25% over 99 min, followed by 25-40% over 5 min, followed by washing | solvent B in A from 4% to 25% over 89 min, followed by 25-40% over 15 min, followed by washing |
| MS1 mode | Positive ion, profile mode on the Orbitrap |  |
| MS1 m/z range | 300-1500 m/z | 350-1400 m/z |
| MS1 resolution | 120,000 | 70,000 |
| MS1 | Max. fill time of 250 ms or AGC target of 50% | Max. fill time of 100 ms or AGC target of 3e6 ions |
| MS2 | Precursor charge: 2-7 |  |
| MS2 isolation window | 1.2 m/z | 1.7 m/z,<br>Top20 method |
| MS2 fragmentation | 30% HCD collision energy | 26 NCE (normalized collision energy) |
| MS2 readout | IonTrap |  |
| MS2 settings | max fill time: 50 ms / 200% normalized AGC target | max fill time: 50 ms / AGC target: 1e5 |
| Dynamic exclusion | 45 s | 30 s |

### Data analysis

#### Processing of raw files

Mass spectrometry raw files were processed using IsobarQuant^53^ and peptide and protein identification was obtained with Mascot 2.5.1 (Matrix Science) using a reference mouse proteome (Uniprot Proteome ID: UP000000589, downloaded May 14, 2016) or human proteome (Uniprot Proteome ID: UP000005640, downloaded June 9, 2020) modified to include known common contaminants and reversed protein sequences. Mascot search parameters were: trypsin; max. 2 missed cleavages; peptide tolerance 10 ppm; MS/MS tolerance 0.02 Da; fixed modifications: Carbamidomethyl (C); variable modifications: Acetyl (Protein N-term), Oxidation (M), SILAC heavy Arg10, Lys8.

#### Data filtering, preprocessing, mapping of gene names, and reproducibility analysis

IsobarQuant output data was analyzed on a protein level in R (https://www.R-project.org) using an in-house data analysis pipeline. In brief, protein data (28,815 unique gene name entries, or combinations of gene names for ambiguous cases across both species and all samples) was filtered to remove contaminants and decoy entries (removed 5,982) and proteins with less than 2 unique quantified peptide matches (removed 6,969), after which gene names were mapped across species from human to mouse using the human and mouse homology classes from the Jackson laboratory (https://www.informatics.jax.org/downloads/reports/HOM_MouseHumanSequence.rpt, accessed June 17, 2020). To enable mapping results across species, all subsequent analysis was done on the gene name level using the corresponding human gene name (9,717 gene name entries in total). For analyzing the reproducibility of the proteomic composition of the independent replicates, signal sums (heavy + light channel) were calculated for each protein in each sample, then normalized using vsn normalization^54^ across all samples. Vsn-normalized intensities were then correlated across all samples using the base R cor() function and visualized using the corrplot() function from the corrplot R package using hierarchical clustering. After this, proteins without a SILAC heavy-to-light (H/L) ratio (i.e., cases where one of the two SILAC channels was not quantified) were also filtered out (removed 1,318 gene name entries, leaving 8,399 gene names in total).

#### Protein copy number estimation

We estimated protein copy numbers per cell using the proteomic ruler method^29^. Briefly, a histone signal was then calculated by summing all histone vsn-normalized signal sums. Copy numbers per cell were then estimated for each protein (on a gene name level) using the following equation:

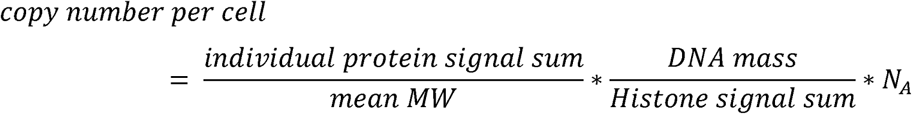

where *N*_A_ is Avogadro’s number, 6.022E23, and the DNA mass is the estimated total signal mass of genomic DNA in each cell (6.5E-12 g for human, and 5.9E-12 g for mouse, assuming 3.099 and 2.728 megabase pairs for human and mouse respectively and mass of a base pair of 615.9 Da^29^), and mean MW is the mean molecular weight of the protein (in case of multiple MWs mapped to a single gene name).

#### Fitting of protein degradation rates (half-lives)

Protein degradation rates (*k*_deg_) were determined as previously described^27,28^. In brief, a linear fit was used to fit ln(H/L+1) over time with the y-intercept fixed to zero. Turnover fits were done only for cases with data in at least two time points in a given replicate (removing 1,131 gene names), yielding estimates for 7,268 gene names. Assuming exponential degradation of all proteins and doubling of the total protein amount during one cell cycle, the slope of this line is defined by *k*_deg_ + *t*_cc_, where *k*_deg_ is the degradation rate constant, and *t*_cc_ is the cell cycle (i.e., doubling) time. We estimated *t*_cc_ from the distribution of all positive slopes of each replicate for each species, assuming that proteins with the 1% lowest slopes turn over only by dilution (*k*_deg_ = 0) and that the slope is thus defined by *t*_cc_. Protein degradation rates were then calculated by correcting for the cell cycle (*k*_deg_ = slope - *t*_cc_). Protein degradation rates were mapped to gene name level by taking the median degradation rate across all protein Uniprot IDs under one gene name, and half-lives were calculated by taking half-life *T*_1/2_ = ln2 / *k*_deg_. For comparisons across species, only gene names with half-lives in at least two biological replicates were considered, and half-lives were consolidated by taking the median across replicates for each species (except for the mouse_2DG sample, where only a single biological replicate was available) (removing 1,133 gene names, leaving a total of 6,135 gene names). The number of overlapping non-negative half-lives (after cell-cycle time correction) across species from these are shown in the respective figures.

#### Statistical comparison of individual protein turnover

To find proteins statistically significantly differing in degradation rates across samples (human v mouse or mouse v mouse_2DG), a comparative fitting approach was used as previously described^31^. Gene name-mapped (see above) H/L ratios were first corrected for *t*_cc_ to enable pooling all data from biological replicates as follows.

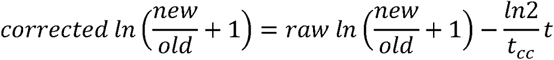

Only gene names with data in at least 3 time points per sample, and a total of at least 8 data points (for human v mouse) or 6 data points (for mouse v mouse_2DG) were included. An F-statistic was calculated for fitting a linear fit to either data from both species together (H0 model) or to each separately (H1 model) (Supplementary Fig. 2e). To account for the heteroscedasticity of the data, the resulting F-statistic was calibrated as described previously^30^ by estimating the “effective degrees of freedom” (d□, d□) from a null dataset, where the species labels were randomized (thus reducing any differences between the two species to those occurring by chance). d□, d□ were estimated for four separate bins of the data, depending on the number of data points available for the comparison. The calibrated F-statistic distribution across both fractions was then used to calculate p-values for each peptide using the pf() function from the stats package in R and corrected for multiple testing using Benjamini-Hochberg correction using the p.adjust() function also from the stats package. Cases with adjusted p-value <= 0.001 were considered as being significantly different in degradation rates across samples tested.

#### Gene set enrichment analysis

GO terms were taken from Uniprot (downloaded from Uniprot on Dec 13, 2022) using the human gene names in our dataset and mapping to the first gene name in Uniprot. All GO term classes (Cellular component, CC; Biological process, BP; and Molecular function, MF) were considered separately. Gene set enrichment analysis was done using the list of gene names ranked according to the difference in degradation rates in human v mouse (or mouse v mouse_2DG) and the fgsea() function from the fgsea package^55^ with the following parameters: minimum set size: 15, maximum set size: 500. Only pathways with an adjusted p-value of < 0.01 as given by fgsea are shown in the figures.

#### Protein subcellular localization

Annotation was done based on GO cellular component terms using the following subset of GO terms: GO:0005576 (extracellular region), GO:0031012 (extracellular matrix), GO:0005886 (plasma membrane), GO:0005856 (cytoskeleton), GO:0005737 (cytoplasm), GO:0005829 (cytosol), GO:0005739 (mitochondrion), GO:0005777 (peroxisome), GO:0005764 (lysosome), GO:0005783 (ER), GO:0005794 (Golgi apparatus), GO:0005634 (nucleus), GO:0031965 (nuclear membrane), GO:0005654 (nucleoplasm), GO:0005730 (nucleolus). Proteins annotated with multiple of these terms were included in all of the annotated groups. Groups were compared with t test using the stat_compare_means() function from the ggpubr package.

#### Data visualization

Figures were prepared using in-house R scripts based mainly on the ggplot2 and ggpointdensity packages. All box plots consist of the median line, box: upper and lower quartiles, whiskers: 1.5 times interquartile range, and points: outliers.

### HES7 oscillation assay

Mouse EpiSCs stably carrying the mHes7 promoter-Fluc-NLS-PEST-UTR reporter and human iPSCs stably carrying the hHES7 promoter-Fluc-NLS-PEST-UTR reporter were differentiated into PSM (mouse: standard protocol; human: 1-step protocol). After induction, the medium was replaced with SCDF medium, in which the concentration of CHIR99021 was reduced to 1 μM, supplemented with D-luciferin. For perturbation experiments, mouse iPSM cells were pre-treated with 2DG for 2 h before medium replacement and maintained in SCDF containing the same concentration of 2DG throughout the measurement, while human iPSM cells were treated with bortezomib at the time of medium replacement. Luminescence from the entire culture dish was continuously recorded using a Kronos Dio Luminometer (Atto).

Traces were analyzed with pyBOAT 0.9.12^56^. For mouse data, a threshold of 300 min was applied for both Sinc detrending and amplitude normalization; for human data, thresholds of 500 min and 300 min were used, respectively. A Fourier-based estimate of the wavelet analysis was used to obtain the distribution of periods and their corresponding power, and the period with maximal power was assigned for each trace.

### Metabolic rate measurement

Metabolic assays were carried out using a Seahorse XF Analyzer (Agilent). iPSM cells were seeded on fibronectin-coated Seahorse plates (Agilent) at a density of 9.1 × 10□ cells/cm² in 100 μl of Seahorse XF DMEM (Agilent) supplemented with glucose (20 mM), pyruvate (1 mM), and glutamine (2 mM). Cells were allowed to attach at room temperature for 10 min and then placed in a 37 °C incubator without CO□ for another 10 min. Subsequently, 400 μl of pre-warmed Seahorse XF DMEM was gently added to each well, resulting in a final volume of 500 μl, taking care not to dislodge the cells. Plates were then incubated at 37 °C without CO□ for an additional 30 min. For the real-time ATP rate assay (Agilent), oligomycin (1 μM), rotenone (0.5 μM), and antimycin A (0.5 μM) were sequentially applied. Samples were also treated with varying concentrations of 2DG. Data were analyzed using the Wave Desktop software and the associated online application provided by the manufacturer.

### Degradation assay

Mouse EpiSCs and human iPSCs stably expressing either the rTetOne promoter-mHes7 (w/o intron)-Nluc construct or the individual localization reporters were used. iPSM differentiation was carried out in the presence of doxycycline (Dox; 100 ng/ml). Expression of Nluc-fusion proteins was initiated by removing Dox and replacing the medium with CDMi supplemented with protected furimazine (Promega). After 4-5 h, once Nluc signals were detected, transcription was stopped by re-adding Dox (100 ng/ml) at time 0, and the decay of Nluc luminescence was monitored with a Kronos Dio luminometer. Protein half-life was estimated by calculating the slope of the log2-transformed signal. A RANSAC algorithm (scikit-learn) was applied to identify the most linear segment of the decay curve. For conditions showing complex degradation dynamics, the threshold parameter of the RANSAC algorithm was reduced to achieve a better overall fit.

### Real-time monitoring of PSM differentiation timing

Human iPSCs stably expressing the hHES7 promoter-Fluc-NLS-PEST-UTR reporter were used to monitor PSM differentiation. At the onset of induction, cells were cultured in SCDF medium in the presence or absence of 2DG. Luciferase activity from the reporter construct was continuously monitored in real time using the same luminescence recording system as described for the oscillation assays (Kronos Dio Luminometer, Atto). Luminescence signals were recorded from the whole culture dish throughout the differentiation process, and traces were analyzed to evaluate the effect of 2DG treatment on the timing of PSM differentiation.

### Quantitative RT-PCR (qRT-PCR) analysis

Total RNA was extracted from human iPSCs differentiating into iPSM using the RNeasy Mini Kit (Qiagen) according to the manufacturer’s instructions. Samples were collected every day during the PSM induction period from both the control and 2DG-treated groups. cDNA was synthesized from 1 µg of total RNA using the PrimeScript™ 1st strand cDNA Synthesis Kit (Takara) with oligo(dT) primers. Quantitative PCR was performed using the LightCycler® 480 SYBR Green I Master (Roche) on a LightCycler® 480 Instrument II (Roche).

mRNA levels were normalized to GAPDH, and relative expression was calculated using the LightCycler® 480 Software (Roche). Target genes included HES7 (for confirmation of reporter readout) as well as other marker genes to evaluate cellular differentiation timing. Primer sequences are listed in Supplementary Table 2.

### Measurements

All measurements were taken from distinct samples (biological replicates) unless stated otherwise.

## Data availability

The mass spectrometry proteomics data have been deposited in the ProteomeXchange Consortium via the PRIDE^57^ partner repository with the dataset identifier PXD051227. Source data are provided as a Source data file.

## Code availability

The custom scripts used are available from Github [https://github.com/mebisuya/SILAC].

## Acknowledgments

We are grateful to K. Stapornwongkul, T. Rayon, and J. Briscoe for their comments. This work was supported by EMBL; the Deutsche Forschungsgemeinschaft (DFG, German Research Foundation) under Germany’s Excellence Strategy - EXC 2068 - 390729961 - Cluster of Excellence Physics of Life of TU Dresden; the European Research Council (ERC) under the European Union’s Horizon 2020 research and innovation program (grant agreement No. 101002564) (to M.E.); PRESTO (grant number JP20332265) from Japan Science and Technology Agency (JST) (to M.M.); the Boehringer Ingelheim Fonds (BIF) PhD fellowship (to J.L.); H.M.H. was supported by a fellowship from the EMBL Interdisciplinary Postdoctoral (EI3POD) Programme under Marie Skłodowska-Curie Actions COFUND (grant number 664726); M.M.S. is supported by the Allen Distinguished Investigator award through the Paul G. Allen Frontiers Group; M.E. is supported by the Alexander von Humboldt Foundation in the framework of the Alexander von Humboldt Professorship endowed by the Federal Ministry of Education and Research.

The human iPSC and mouse EpiSC lines were provided by the RIKEN BRC through the National BioResource Project of the MEXT, Japan.

## Author contributions

M.M. and M.E. designed the work and wrote the manuscript. H.M.H. and M.M.S performed and analyzed SILAC-proteomics. M.M. prepared the SILAC samples and performed the other experiments. M.M. and J.L. analyzed the non-SILAC data. All authors contributed to the manuscript and approved the final version.

---

**Supplementary Figure 1 Protein copy number a,** Protein copy numbers per cell in mouse and human iPSM cells as estimated using the proteomic ruler method (see Methods). N = 6715 (mouse), 6940 (human). P-value is from two-sided Mann-Whitney U test. Box plot consists of median line, box: upper and lower quartiles, whiskers: 1.5 times interquartile range, points: outliers. **b,** Scatter plot comparing protein copy numbers per cell measured in human and mouse iPSM. N = 5345.

**Supplementary Figure 2 SILAC data analysis a**, Principle of dynamic SILAC. The ratio of new (in this case peptides carrying heavy-labeled amino acids) to old (light) peptides was measured on the MS1 level on the mass spectrometer. **b**, Equations describing the change in new and old proteins assuming a steadily-growing (cell cycle time *t*_cc_) but otherwise steady-state proteome. **c,** The first equation in b defines a straight line of ln(new/old + 1) over time with slope *k*_deg_ + ln2/*t*_cc_. *k*_deg_: degradation rate constant. **d**, To compare degradation rates across samples, the measured signal was first corrected for *t*_cc_, yielding a straight line with slope *k*_deg_. **e**, Comparative fitting approach. For each protein, it was estimated whether the data from two samples (e.g., mouse and human iPSM cells) were better fitted by two separate linear fits (H1 model), or whether a single fit was sufficient (H0 model). **f**, Using the residuals from the fits in e, an F-statistic was calculated to estimate the increase in goodness of fit when going from H0 to H1. For the estimation of the term d_2_/d_1_, see Methods and references therein.

**Supplementary Figure 3 Cell cycle time estimation and reproducibility of SILAC samples a,** Cell cycle time was estimated by using the top 1% of the slowest degradation rates for each replicate. Graph indicates mean ± SD. **b,** Number of unique gene names per sample before filtering for SILAC quantification (total intensity, heavy + light). **c,** Pairwise correlation across samples based on vsn-normalized total intensity. Samples were clustered using hierarchical clustering. **d,** Scatterplots comparing the replicates of SILAC samples based on half-lives. **e,** Fold changes between human and mouse half-lives were evenly distributed across the median protein abundance of human and mouse. The color code is based on Fig. 1d. N = 3630.

**Supplementary Figure 4 Degradation assays of localization reporters a,** Schematic representation of the localization reporter constructs. rTetOne: reverse TetOne; aa: amino acids; Nluc: Nano luciferase. The coding sequence (CDS) and 3’ untranslated region (UTR) of human HES7 gene were added to destabilize the construct mRNA. **b,** Degradation assays of localization reporters in human and mouse iPSM. The transcription of the localization reporters was halted by Dox at time 0, and the decay of the luciferase activity was monitored. **c,** Degradation assays of localization reporters in the presence of a lysosomal inhibitor bafilomycin A1. Mouse iPSM was treated with 80 nM bafilomycin from 4 h before Dox addition. Control (mouse) data are the same as b. **d,** Degradation assays of the nuclear localization reporter in the presence of a proteasome inhibitor MG-132. Mouse iPSM was treated with 10 µM MG-132 together with Dox at time 0. **b-d,** Shading indicates mean ± SD (N = 3).

**Supplementary Figure 5 Half-life estimation in human and mouse iPSM cells** Fitting of protein degradation shown in Fig. 2c and Supplementary Fig. 4b. Dashed lines indicate the most linear region considered by the RANSAC algorithm for the fitting. Slope of the fitted line is shown, and it was converted to the half-life using the equation: half-life = -1/slope.

**Supplementary Figure 6 Effects of glycolysis inhibition on the segmentation clock and protein degradation a,** Dose-dependent effect of a glycolysis inhibitor 2-Deoxy-D-glucose (2DG) on the segmentation clock period. Mouse iPSM was pre-treated with 2DG from 2 h before time 0, and the oscillatory activity of the Hes7 promoter-luciferase reporter was monitored. **b,** Hes7 oscillation periods estimated from a. **c,** Dose-dependent effect of 2DG on Hes7 protein degradation. Mouse iPSM was pre-treated with 2DG from 4 h before Dox addition. The transcription of Hes7-Nluc was halted by Dox at time 0, and the decay of the luciferase activity was monitored. **d,** Hes7 protein half-life calculated from c. **a,c,** Shading indicates mean ± SD (N = 3). **b,d,** Graphs indicate mean ± SD (N = 3). P-values are from two-sided Dunnett’s test against the indicated controls. Human data is from Matsuda et al, Science (2020)^6^.

**Supplementary Figure 7 Effects of glycolysis inhibition on metabolic rates** Mouse iPSM was treated with different concentrations of 2DG and measured by the Seahorse analyzer. Graphs indicate mean ± SD (N = 3). **a,** Proton efflux rate (PER). **b,** Oxygen consumption rate (OCR). Oligomycin (Oligo) and Rotenone + Antimycin A (Rot + AA) were added at the marked time points. **c,** Basal (before adding 2DG) and induced (after 2DG) ATP production rates calculated from a,b. Glycolytic ATP production rate (glycoATP), mitochondrial ATP production rate (mitoATP), and total ATP production rate were estimated. **d,** Relative ATP production rates. The ratio of induced rates (after adding 2DG) to basal rates (before 2DG) was calculated from c.

**Supplementary Figure 8 Effects of glycolysis inhibition on protein stability profiles** Fold changes between mouse_2DG and mouse half-lives across the median protein abundance of mouse_2DG and mouse. a.u.: arbitrary units. N = 3741. The color code in b is based on Fig. 3b.

**Supplementary Figure 9 GO analysis a,** GO cellular component (CC), molecular function (MF), and biological process (BP) terms statistically significantly enriched in genes that showed slower degradation in mouse_2DG compared with mouse. **b,** GO BP terms statistically significantly enriched in genes that showed slower degradation in mouse_2DG compared with human. **a,b,** P-values are probability estimates that the given enrichment/depletion of the particular gene sets are non-random (two-sided, see fgsea^55^) adjusted for multiple testing using the Benjamini-Hochberg correction. GO terms with adjusted p-value < 0.01 are shown.

**Supplementary Figure 10 Degradation assays of localization reporters in the presence of 2DG** Degradation assays of the localization reporters in the presence of 2DG. Mouse iPSM was pre-treated with 2DG from 4 h before Dox addition. The transcription of the localization reporters was halted by Dox at time 0, and the decay of the luciferase activity was monitored. Shading indicates mean ± SD (N = 3).

**Supplementary Figure 11 Half-life estimation in mouse iPSM treated with 2DG** Fitting of protein degradation shown in Supplementary Fig. 10. Dashed lines indicate the most linear region considered by the RANSAC algorithm for the fitting. Slope of the fitted line is shown, and it was converted to the half-life using the equation: half-life = - 1/slope.

**Supplementary Figure 12 PSM marker expression during cellular differentiation in the presence of 2DG** qRT-PCR analysis of MSGN1 and TBX6 mRNA expression during PSM differentiation from human iPSCs in the presence of 2DG. Graphs indicate mean ± SD (N = 3). **a,c,** Expression in control and 2DG-treated cells. **b,d,** Expression in 2DG-treated cells only, shown with the same time axis as a,c, but with a rescaled y-axis to improve visibility.

**Supplementary Table 1** Measured half-lives for ∼5000 proteins in mouse, human, and mouse_2DG samples.

**Supplementary Table 2** Primer sequences for qRT-PCR.

