## Supplementary Figures for "Systematic differences in protein stability underlie species-specific developmental tempo"

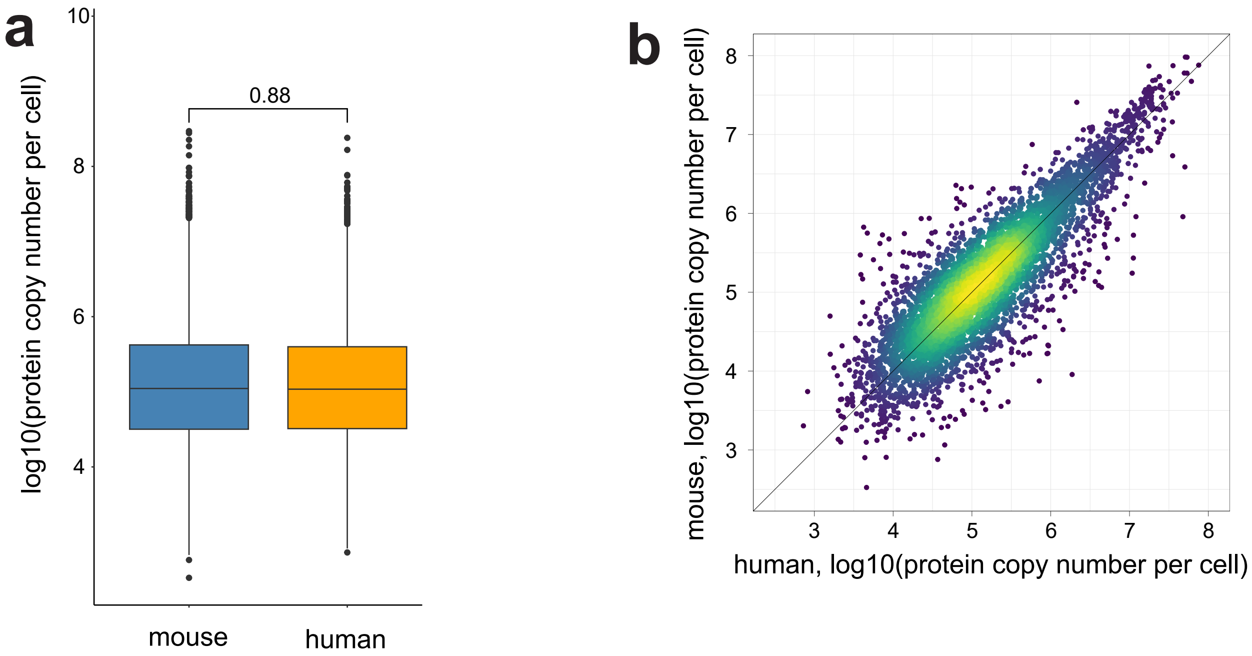


Supplementary Fig. 1

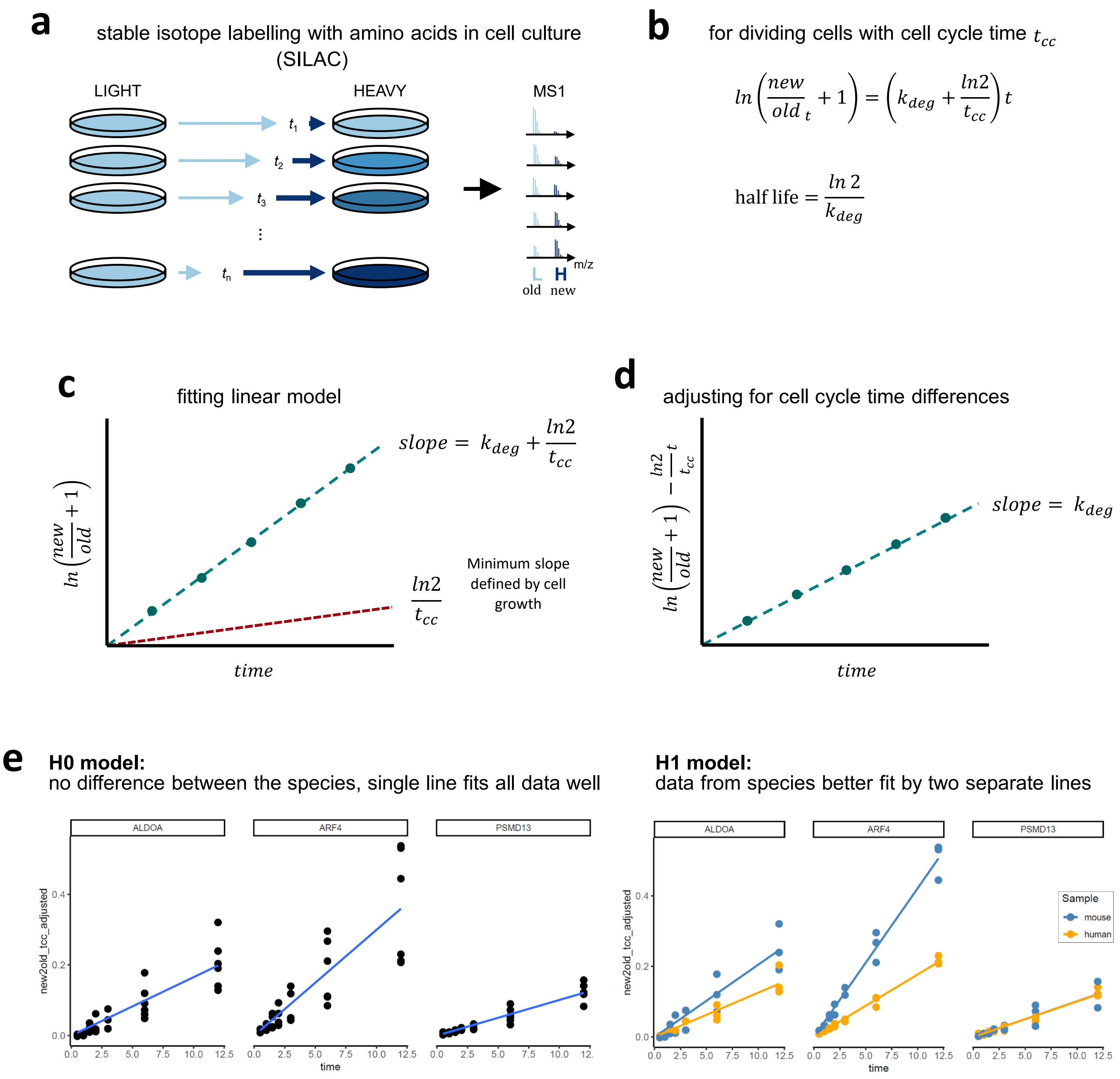


**f**

$$F = \frac{d_2}{d_1}\frac{(RSS_0 - RSS_1)}{RSS_1}$$

$F$ F-statistic

$RSS_0$ residual sum of squares for H0 model

$RSS_1$ residual sum of squares for H1 model

$\frac{d_2}{d_1}$ ratio of degrees of freedom. Here, determined experimentally from data, due to heteroscedasticity

Supplementary Fig. 2

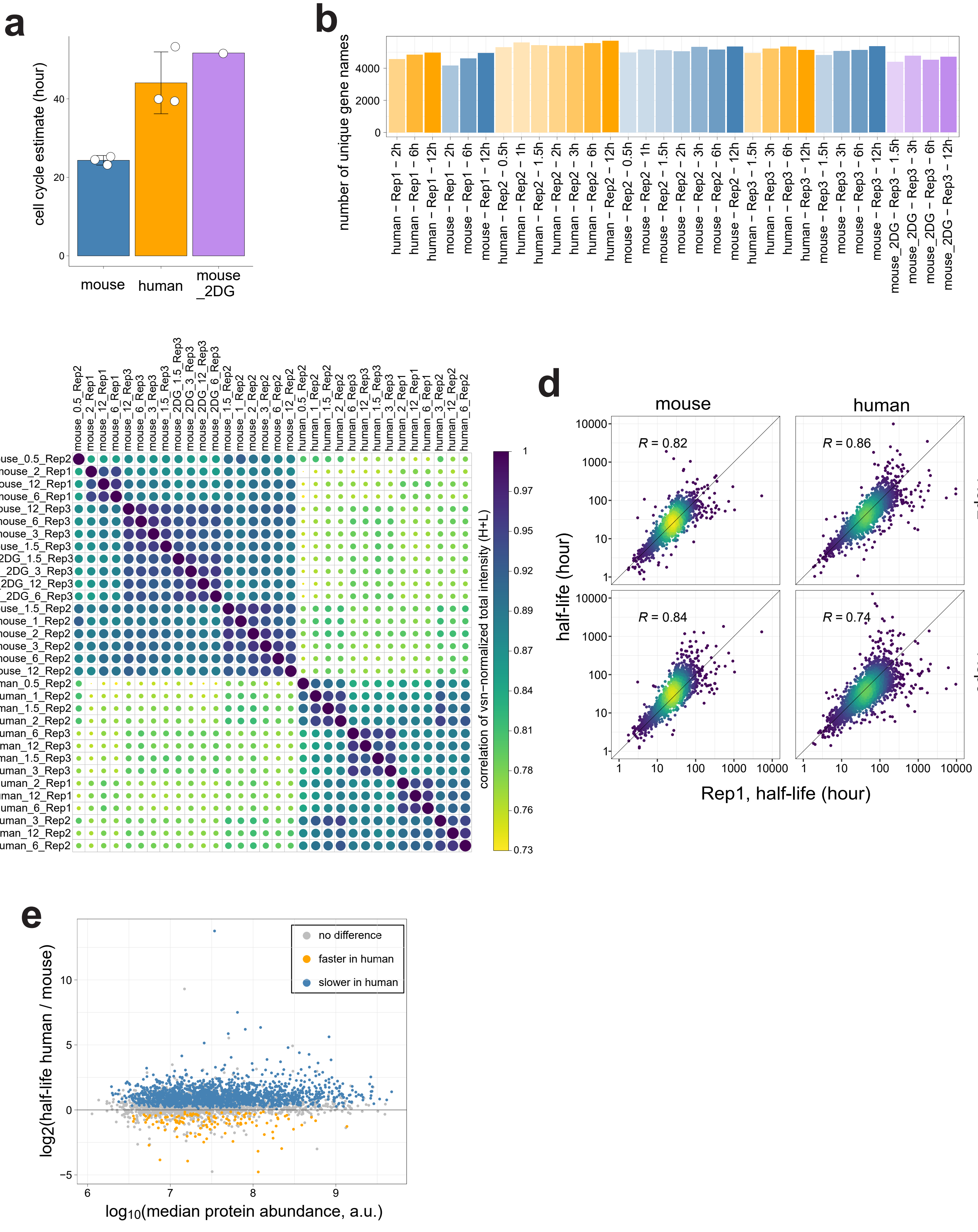


Supplementary Fig. 3

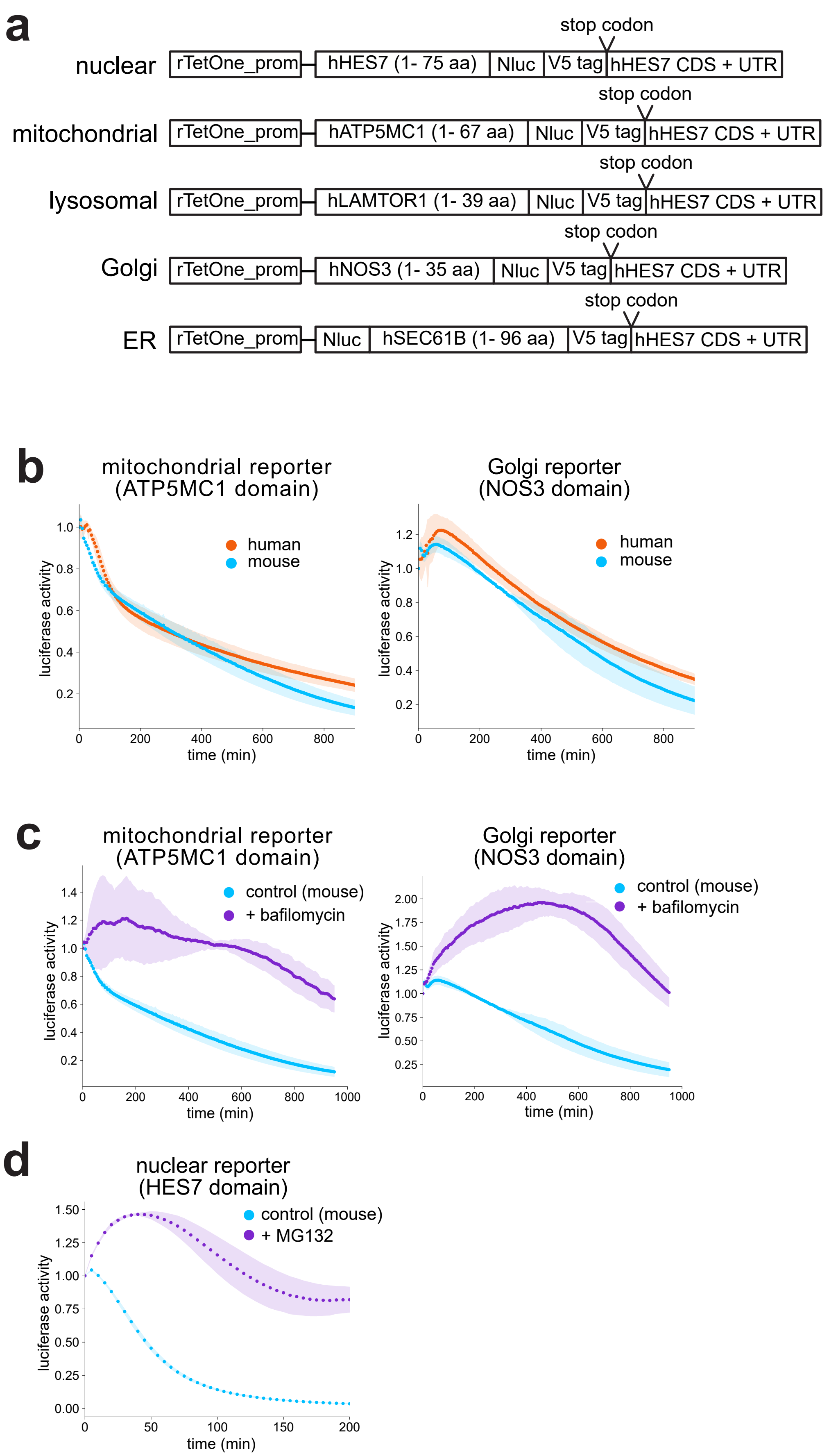


Supplementary Fig. 4

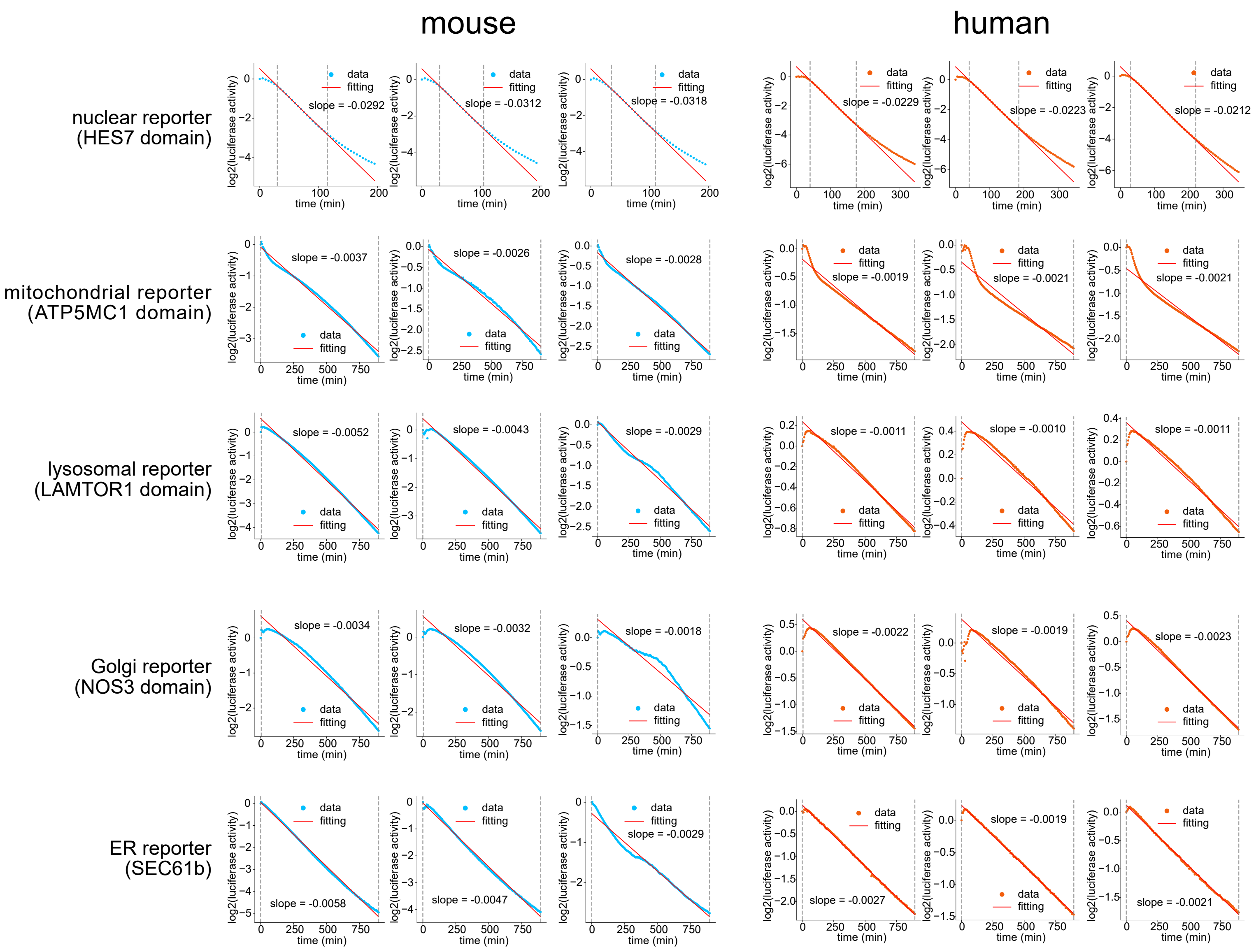


Supplementary Fig. 5

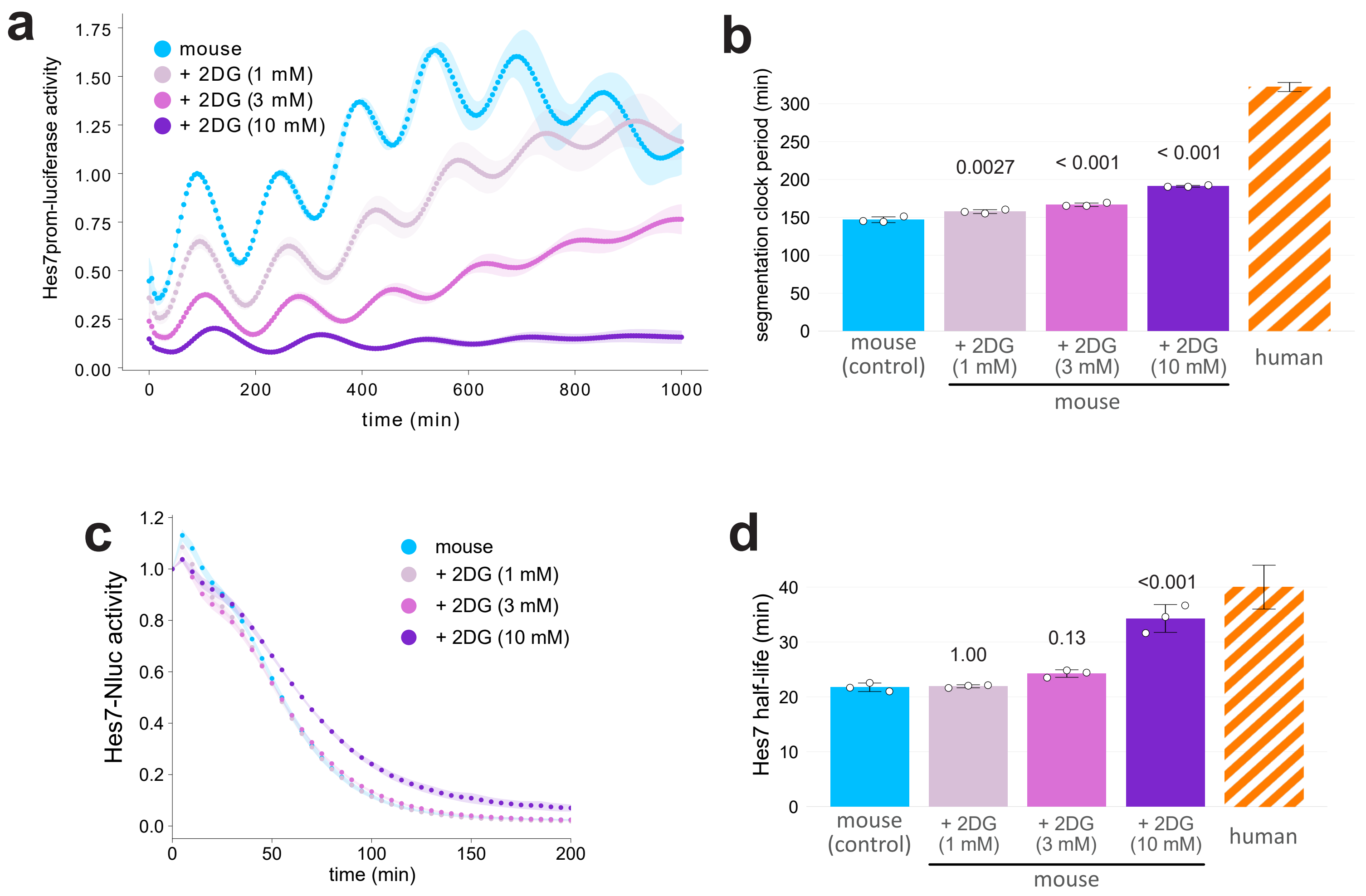


Supplementary Fig. 6

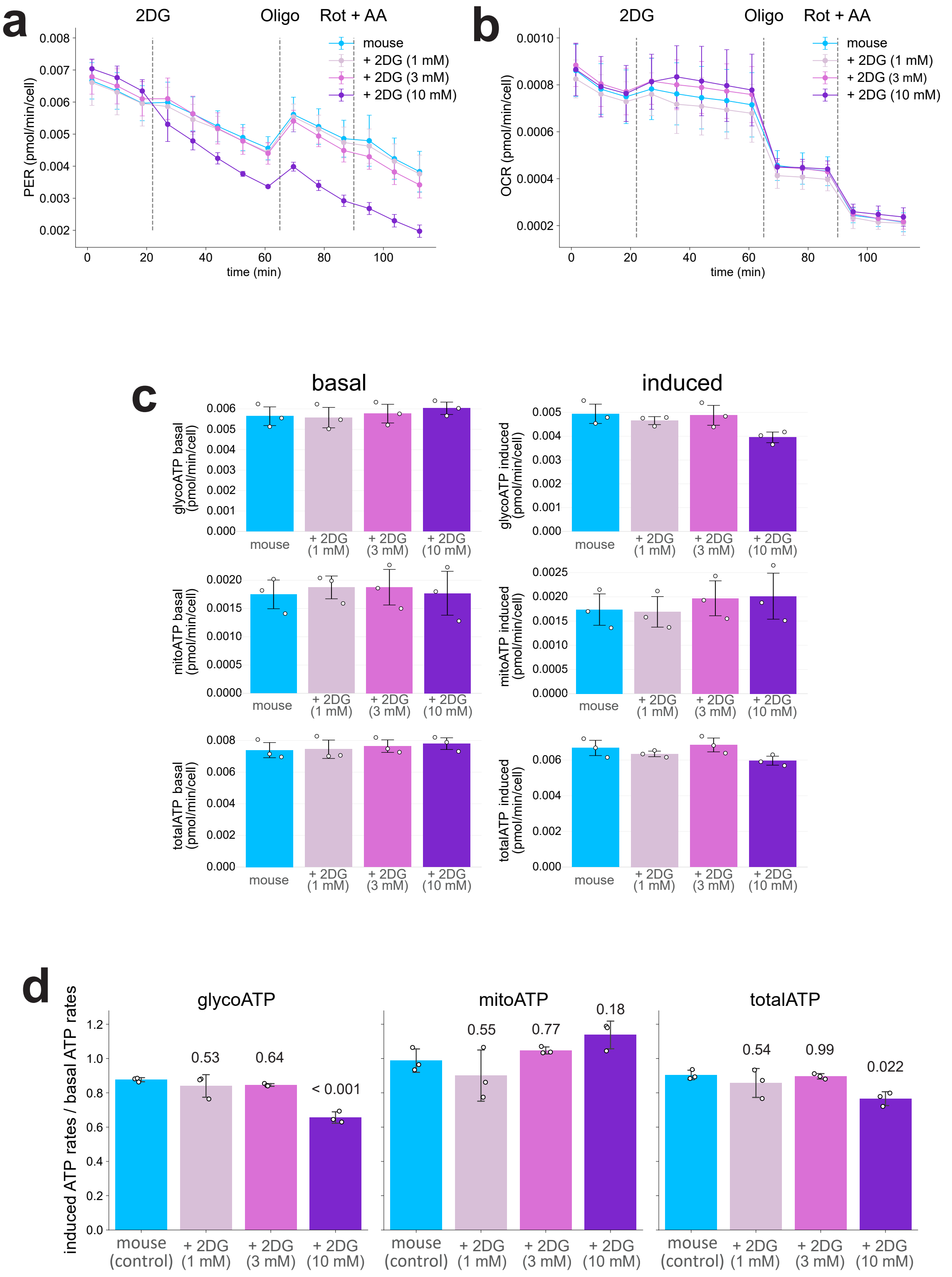


Supplementary Fig. 7

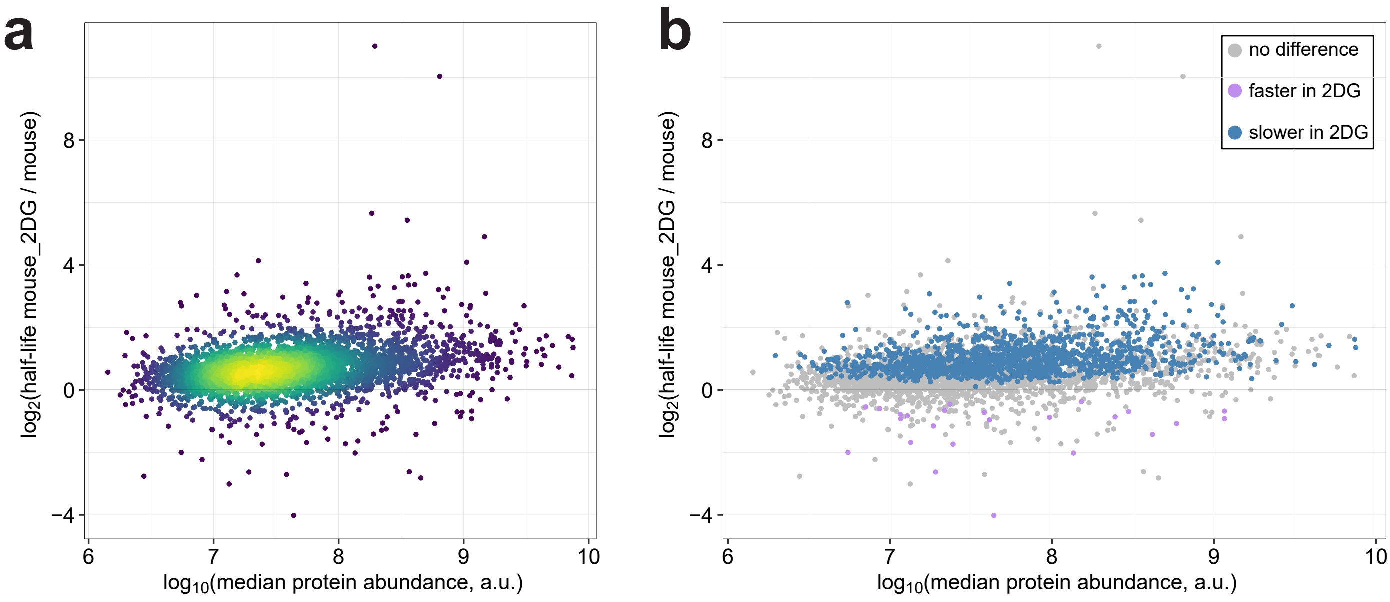


Supplementary Fig. 8

**a**

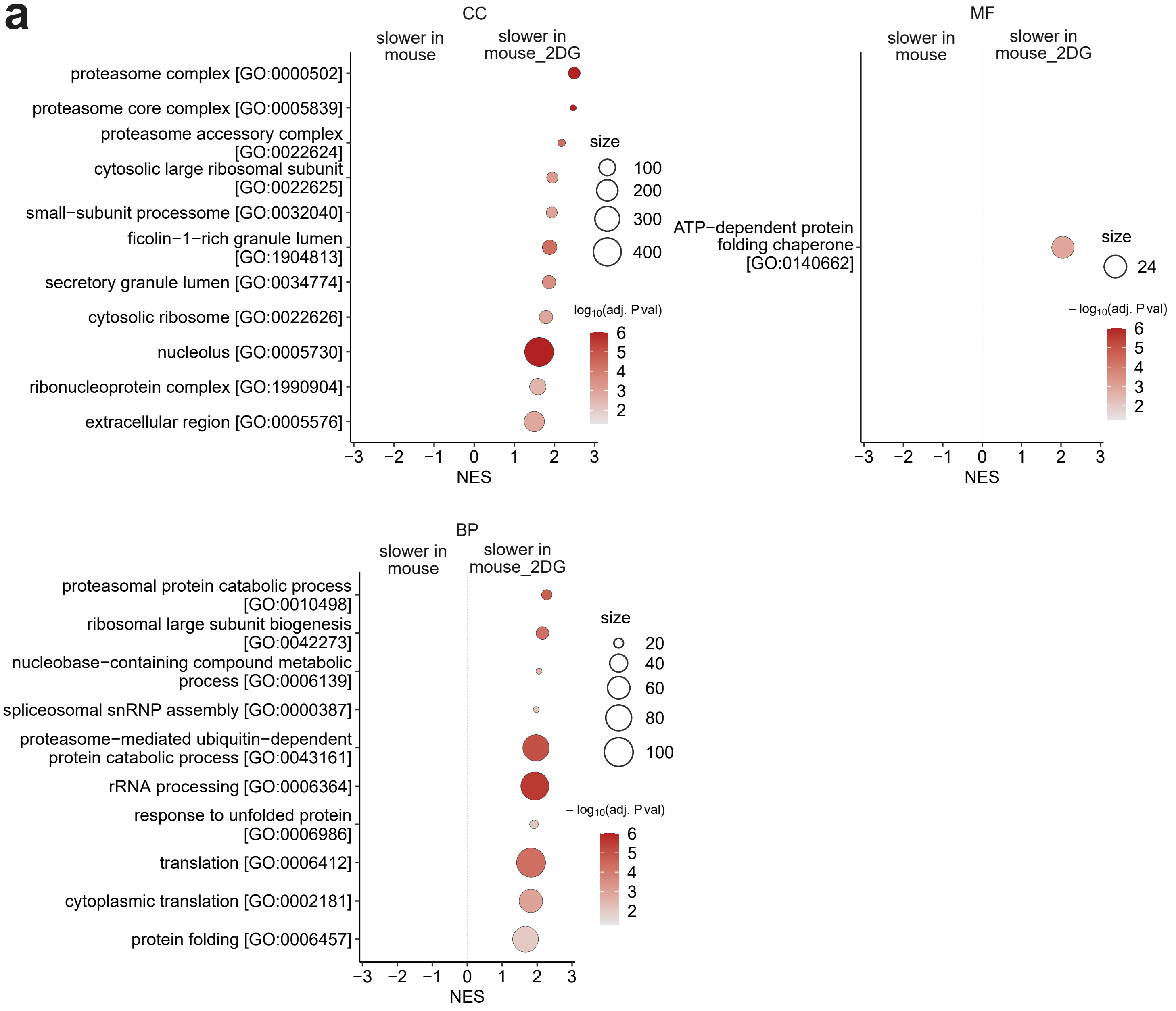


**b**

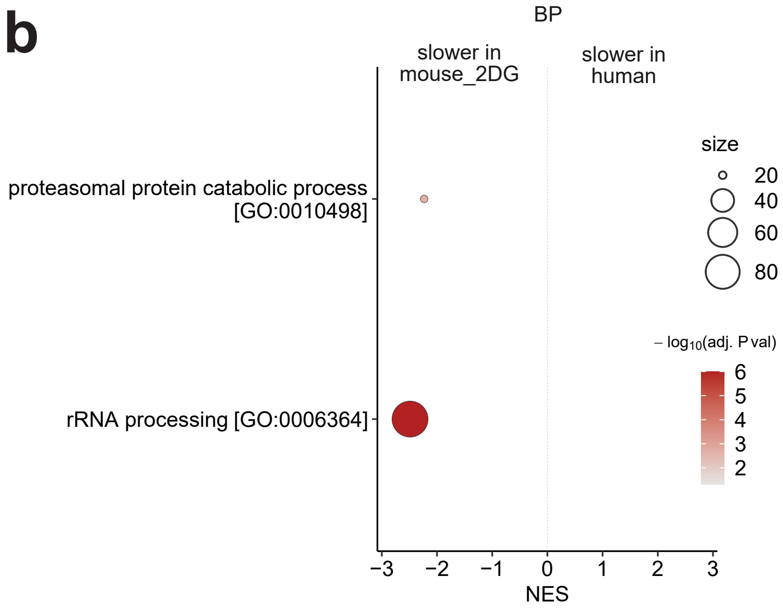


Supplementary Fig. 9

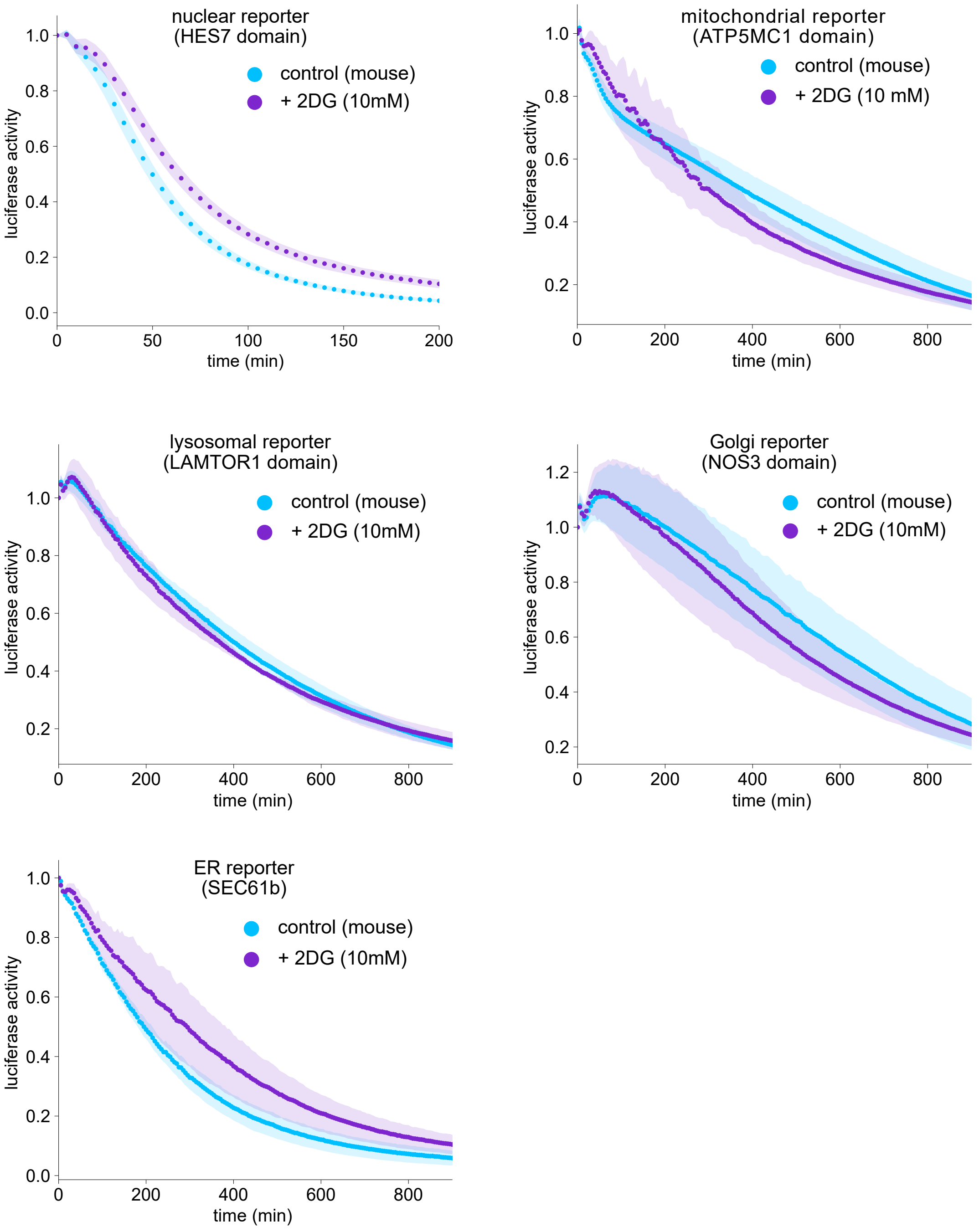


Supplementary Fig. 10

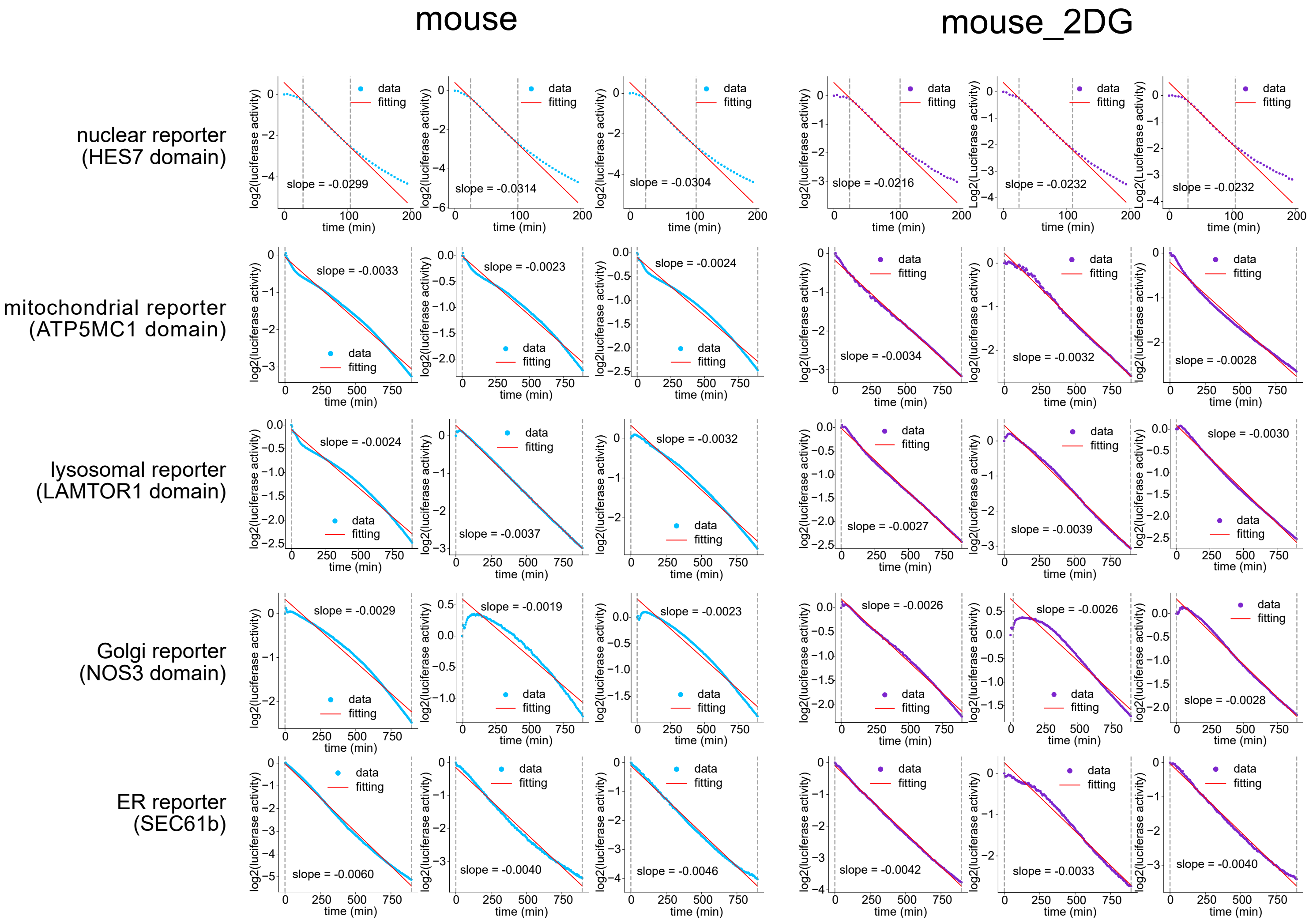


Supplementary Fig. 11

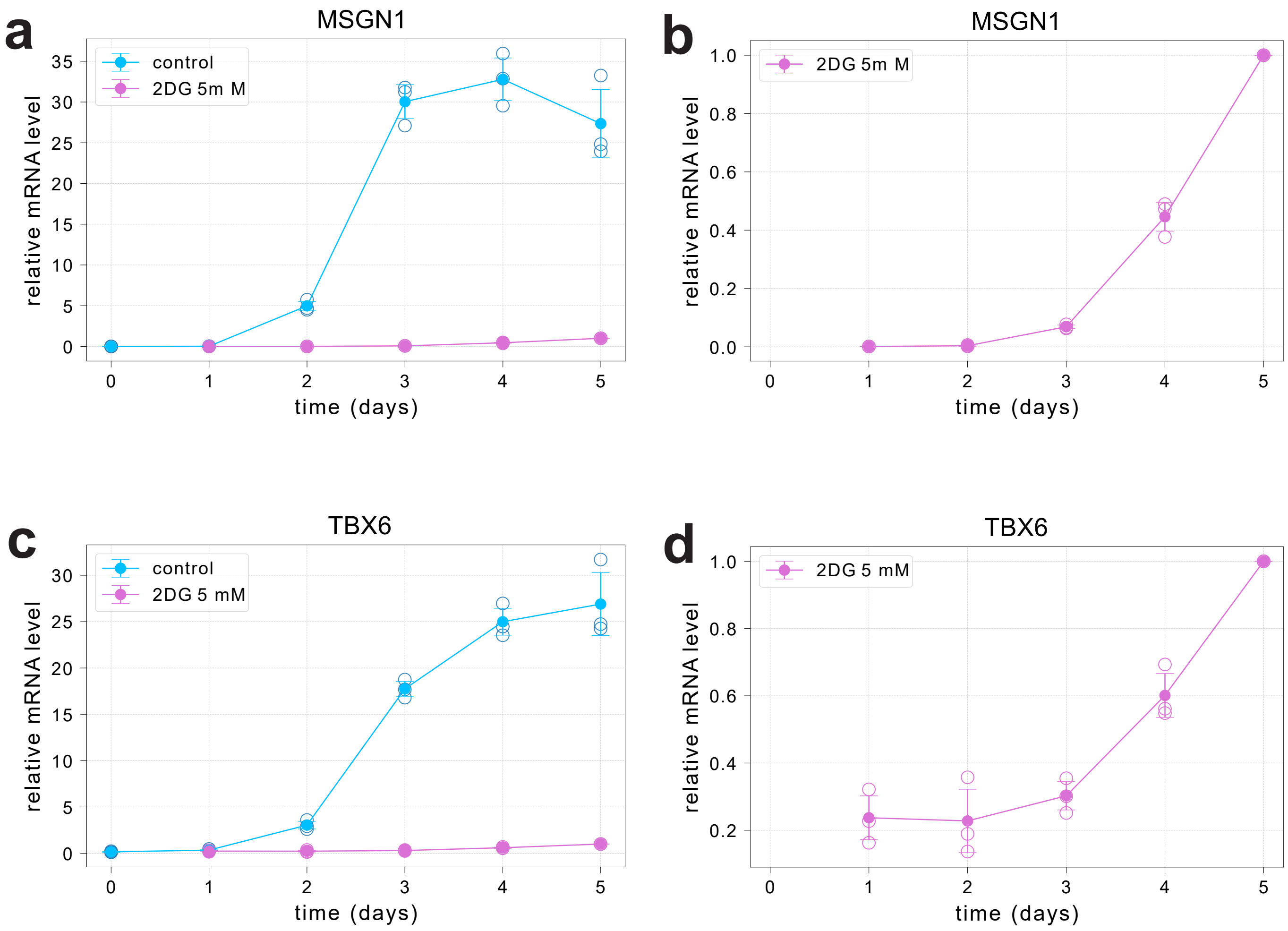


Supplementary Fig. 12
